# Multi-omic lineage tracing predicts the transcriptional, epigenetic and genetic determinants of cancer evolution

**DOI:** 10.1101/2023.06.28.546923

**Authors:** F. Nadalin, M.J. Marzi, M. Pirra Piscazzi, P. Fuentes, S. Procaccia, M. Climent, P. Bonetti, C. Rubolino, B. Giuliani, I. Papatheodorou, J.C. Marioni, F. Nicassio

## Abstract

Cancer is a highly heterogeneous disease, where phenotypically distinct subpopulations coexist and could be primed to different fates. Both genetic and epigenetic factors may drive cancer evolution, however little is known about whether and how such a process is pre-encoded in cancer clones. Using single-cell multi-omic lineage tracing and phenotypic assays, we investigate the predictive features of either tumour initiation or drug tolerance within the same cancer population. Clones primed to tumour initiation *in vivo* display two distinct transcriptional states at the baseline. Remarkably, these states share a distinctive DNA accessibility profile, highlighting an epigenetic basis for tumour initiation. The drug tolerant niche is also largely pre-encoded, but only partially overlaps the tumour-initiating one and evolves following two genetically and transcriptionally distinct trajectories. Our study highlights coexisting genetic, epigenetic and transcriptional determinants of cancer evolution, unravelling the molecular complexity of pre-encoded tumour phenotypes.

## MAIN

Cancer adopts evolutionary pathways that sustain the disease. Aggressive tumour behaviours, such as the dissemination to distant organs, diminished susceptibility to treatment, and disease relapse, result from either selection or adaptation processes, possibly intertwined (Marine et al. 2020). When a selective process occurs, the fate of a cancer clone is determined at the root of the evolutionary process. In this case, the heterogeneity of tumour phenotypes can, at least in principle, be identified ahead of selection (Marusyk et al. 2012). The pre-existence of aggressive phenotypes has been linked to the so-called cancer stem cell (CSC) theory (Visvader and Lindeman 2008) and observed in leukemia (Lapidot et al. 1994, Bonnet and Dick 1997) and solid tumours, such as colon (Ricci-Vitiani et al. 2007) and breast cancer (Al-Hajj et al. 2003) (Batlle and Clevers 2017), as well as glioma (Singh et al. 2004) (Lathia et al. 2015). According to such a model, tumour cells are not all equal, instead a stem-like cancer niche exists that is primed to sustain most of the aggressive phenotypes, such as tumour re-initiation, metastatic dissemination potential, and capacity to survive cytotoxic treatments (Basile and Aplin 2012).

Predicting cancer phenotypes requires linking the molecular state of a clone to its fate with high precision. Without *a priori* information, tumour phylogeny can be inferred from somatic mutations (Ross and Markowetz 2016, McCarthy et al. 2020, Satas et al. 2020, Zhou et al. 2020); however, this approach is limited by the high sparsity of single-cell data. Single-cell lineage tracing consists in inserting barcodes in the genome of the cells with the aim of tracing their progeny (Dixit et al. 2016, Biddy et al. 2018, Weinreb et al. 2020, Jindal et al. 2022). In cancer, this approach has been used to investigate clonality in metastases (Simeonov et al. 2021), survival upon cytotoxic treatment (Gutierrez et al. 2021, Oren et al. 2021), as well as to dissect the clonal origin of primary tumour and metastasis growth (Karras et al. 2022), possibly *in vivo* (Yang et al. 2022). However, these studies mainly focus on the evolutionary trajectories, rather than on the driving molecular features of pre-existing phenotypes.

Tumour evolutionary diversity can have either a genetic or non-genetic origin (Salgia and Kulkarni 2018, Black and McGranahan 2021). Single-cell multi-omics has recently emerged as a promising tool to study cancer evolution (Nam et al. 2021). Here, we combine single-cell multi-omics with lineage tracing in a unique framework, which allows profiling clone, gene expression, and chromatin accessibility information simultaneously at single-cell resolution. Using phenotypic assays on barcoded cells, we identify the clones endowed with aggressive cancer behaviours typical of the stem-like cancer niche, specifically the tumour-initiating capacity and drug tolerance. Subsequently, we extract robust transcriptional, epigenetic, and genetic features of naïve cells and associate them to clonal subpopulations. By integrating these multiple layers of information, we identify the regulatory elements that predict cancer evolution in response to adverse environmental conditions. Finally, by tracing the transcriptional changes of clones across time, we unravel the role of pre-existing molecular features in shaping the differentiation breadth of stem-like subpopulations.

## RESULTS

### SUM159PT exhibit high transcriptional plasticity and comprise three distinct subpopulations: S1, S2 and S3

To investigate the potential of cancer cells to promote tumour initiation and escape cytotoxic treatment, we combined single-cell sequencing with phenotypic assays. We selected SUM159PT, a triple-negative breast cancer cell line (TNBC), as model system. SUM159PT belong to the claudin-low mesenchymal subtype (Prat et al. 2013) and is characterised by i) a nearly diploid genotype, but bearing a specific set of mutations typically associated with TNBC intrinsic heterogeneity [HRAS, PIK3CA, TP54 and MYC amplification (Saunus et al. 2018)] ii) an intrinsic heterogeneity, with an underlying variability in the expression of epithelial and mesenchymal genes and a small proportion of cells in a CSC state (Gupta et al. 2011, Bierie et al. 2017) iii) an aggressive phenotype driven by the CSC component, being highly tumorigenic and invasive *in vivo* (Gupta et al. 2011, Bierie et al. 2017, Watson et al. 2021). A single-cell lineage tracing approach was used to link the molecular state of a cell to its phenotypic fate (**Figure 1A** and **S1A**). To obtain ∼10,000 distinct genetic barcodes (GBC), 100,000 SUM159PT cells were infected with a lentiviral pool at MOI = 0.1 and subsequently FAC-sorted to retain only the transduced fraction (Dixit et al. 2016). Endogenous as well as GBC-carrying transcripts were then captured by single-cell RNA-seq (scRNA-seq). The parental population was sampled and processed at two time points, T0 and T1, separated by 13-15 days (see **Figure 1B**). At basal state, between 5017 and 5996 unique clones were found in the two replicates and between 83% and 88% of high-quality cells were assigned a clone identity (see **Figure S1B** and **Suppl. Table 1**), making the lineage of even rare cell subpopulations accessible to analysis. The distribution of clones at the two time points was similar, highlighting that no spontaneous clone selection occurs in the time frame (see **Figure S1C**). Moreover, 68% and 57% of the clones respectively detected in T0 and T1 were shared between the two time points, with > 50% of clones constituted by a single cell in each sample (see **Figure S1D** and **Suppl. Table 1**). When evaluating the relationship between clonality and gene expression profile at basal state, cells stemming from a common clone at the moment of infection, hereafter *sister cells*, were on average only slightly more similar to one another compared to non-sisters (see **Figure 1C**). We next asked whether the transcriptional similarity between sister cells is clone-specific — in other words, whether some clones show a distinctive gene expression profile and other clones are more plastic [a similar approach is proposed in (Weinreb et al. 2020)]. We detected seven distinct gene expression clusters in T0 and T1, respectively (see **Figure 1D** and **S1E** and **Suppl. Table 1**) and compared the clone content of every cluster pair across the two time points using a *clone sharedness* score (see Methods and **Figure 1E**). Most clones that clustered together in T0 were mapped to multiple distinct clusters in T1, and *vice versa*, suggesting a high transcriptional plasticity already at baseline. In contrast, three cluster pairs in T0 and T1, respectively comprising 28% and 23% of the cells at the two time points, showed mutually high clone sharedness (see Methods and **Figure 1F**); we conclude that these subpopulations are transcriptionally stable and we will refer to them as S1, S2, S3 hereafter. They respectively comprise 3.6%, 14.7%, and 7.4% cells on average. We obtained a gene expression signature for each of them (see **Figure 1G** and **Suppl. Table 2**) that is independent of cell-culture effect. Of note, S1 was enriched in genes involved in collagen processing and matrix remodelling (see **Figure S1F** and **Suppl. Table 3**). S1 cells showed upregulation of S100A4, a gene associated with metastatic behaviours (Fei et al. 2017) (Simeonov et al. 2021, Low et al. 2023), and TM4SF1, whose role in promoting cell proliferation and invasion in epithelial tumours has been assessed (Gao et al. 2016, Xing et al. 2017, Yang et al. 2017, Chen et al. 2022, Hou et al. 2022). The microRNA-205 host gene (MIR205HG) was found as S2-specific and has been associated to basal cells, epithelial-to-mesenchymal transition, and multiple cancer diseases (Dong et al. 2019, Liu et al. 2020). S3 was distinguished by the expression of FEZ1, a microtubule adaptor (Alborghetti et al. 2011), and RPS25, a gene acting on cellular response to stress by downregulating p53 (Zhang et al. 2013). In conclusion, single-cell lineage tracing revealed that SUM159PT exhibit high transcriptional plasticity, but comprise three distinct, transcriptionally stable subpopulations.

**Figure 1.**
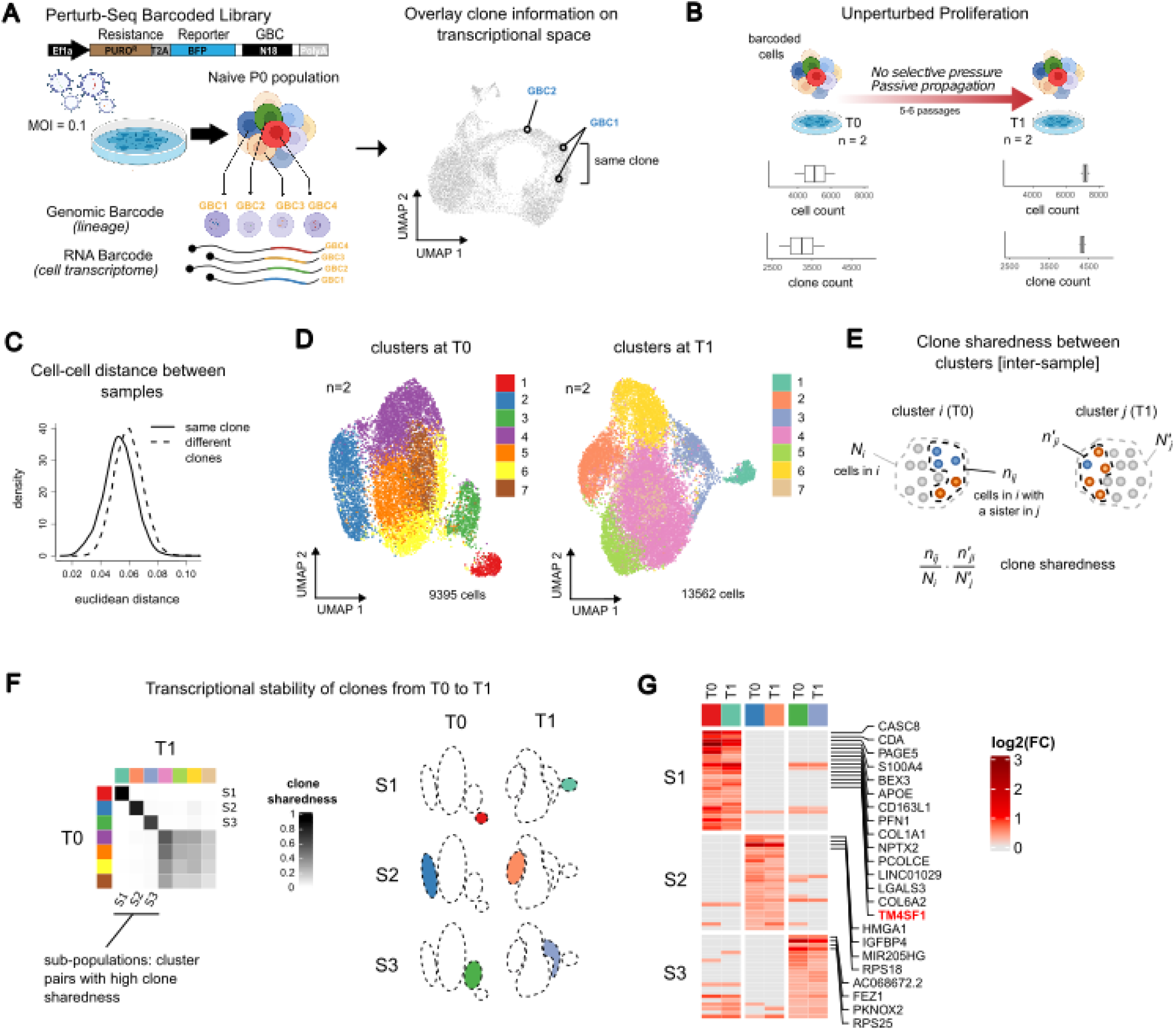
**A.** SUM159PT cells were infected with a lentiviral library of unique barcodes (Perturb-seq GBC library) at 0.1 Multiplicity of Infection (MOI) (estimated by the % of BFP+ cells in FACS). The readout of each cell is its lineage, captured by genetic barcodes (GBCs), and gene expression profile, captured by cDNA reads sequenced at the 3’ end. The final readout (on UMAP representation) is clone information overlaid on the gene expression space at single -cell resolution. **B.** Experimental design, passive propagation. Top: barcoded SUM159PT cells from the same infection experiment were processed by scRNA-Seq at two distinct passages of cell culture (T0 and T1, n=2 replicates per time point). Bottom: number of detected clones and cells assigned to clones across the replicates. **C.** Gaussian kernel density of Euclidean distances between sister cells (solid line) and non-sister cells (dashed line) computed on the integrated space of T0 and T1 (see Methods). **D.** UMAP representation of T0 (9395 cells) and T1 (13562 cells) coloured by cluster; only cells assigned to clones are shown. **E.** Definition of the clone sharedness score between two clusters, *i* and *j*, in T0 and T1. **F.** Clone sharedness scores between T0 and T1. Left: heatmap where rows are clusters in T0 (as in 1D), columns are clusters in T1, and entries are clone sharedness scores for each pair. Rows and columns are sorted according to the pairs with the highest score. The three top-scoring pairs, referred as subpopulations (S1, S2, S3) are shown on UMAP (right panel). **G.** Subpopulation signatures. Heatmap where rows are genes, columns are subpopulations split by time point, and entries are log_2_(FC) values between a subpopulation and its complement at the same time point. Columns are annotated with the same colour as in **D**. and **E**. The 25 genes in the subpopulation gene signature (see Methods) showing highest log_2_(FC) in T0 are shown. The top 15 (S1) or top 4 (S2, S3) are labelled. The surface marker TM4SF1, highlighted in red, is used for sorting the S1 subpopulation hereafter.

### Cancer clones promote tumour initiation in a non-stochastic manner

To investigate the tumour-initiating capacity of SUM159PT, we transplanted barcode-labelled cells into the mammary fat pads of nine NSG (NOD/SCID/IL2Rγ_c_^-/-^) immunodeficient mice and then evaluated the barcode composition in each primary tumour. We isolated tumour cells and extracted the genomic DNA (gDNA), which was then amplified and sequenced (**Figure 2A**). Noteworthy, the GBC count measured in bulk (parental cells) recapitulates the actual clone abundance, measured as the relative number of cells per clone in single-cell samples (**Figure S2A-B**). Only between 3% and 33% of SUM159PT clones contributed to tumour formation (**Figure 2B** and **Suppl. Table 4**), showing a heterogeneous yet deep clone selection. The size of clonal subpopulations greatly varied within a tumour; on average, the top 1% abundant clones covered more than 50% of the entire tumour mass (**Suppl. Table 4**), and this was not merely a consequence of higher initial abundance (see below). This picture suggests a variable tumour initiation potential among surviving clones *in vivo*. In epithelial cancers, the tumour initiation potential has been regarded as an intrinsic feature of cells, rather than a feature that is acquired during tumour formation (Batlle and Clevers 2017). Consistently, we observed that a limited set of clones was recurrent and covered a high proportion of the tumour mass compared to sporadic ones (**Figure 2C**). In particular, 138 clones were significantly more abundant in at least 6 out of 9 tumours compared to their average abundance at basal state (Fisher’s exact test, see Methods). We refer to these as *tumour initiating clones* (TICs) hereafter (**Figure 2D**). We conclude that the tumour-initiating capacity of SUM159PT cells is largely pre-encoded.

**Figure 2.**
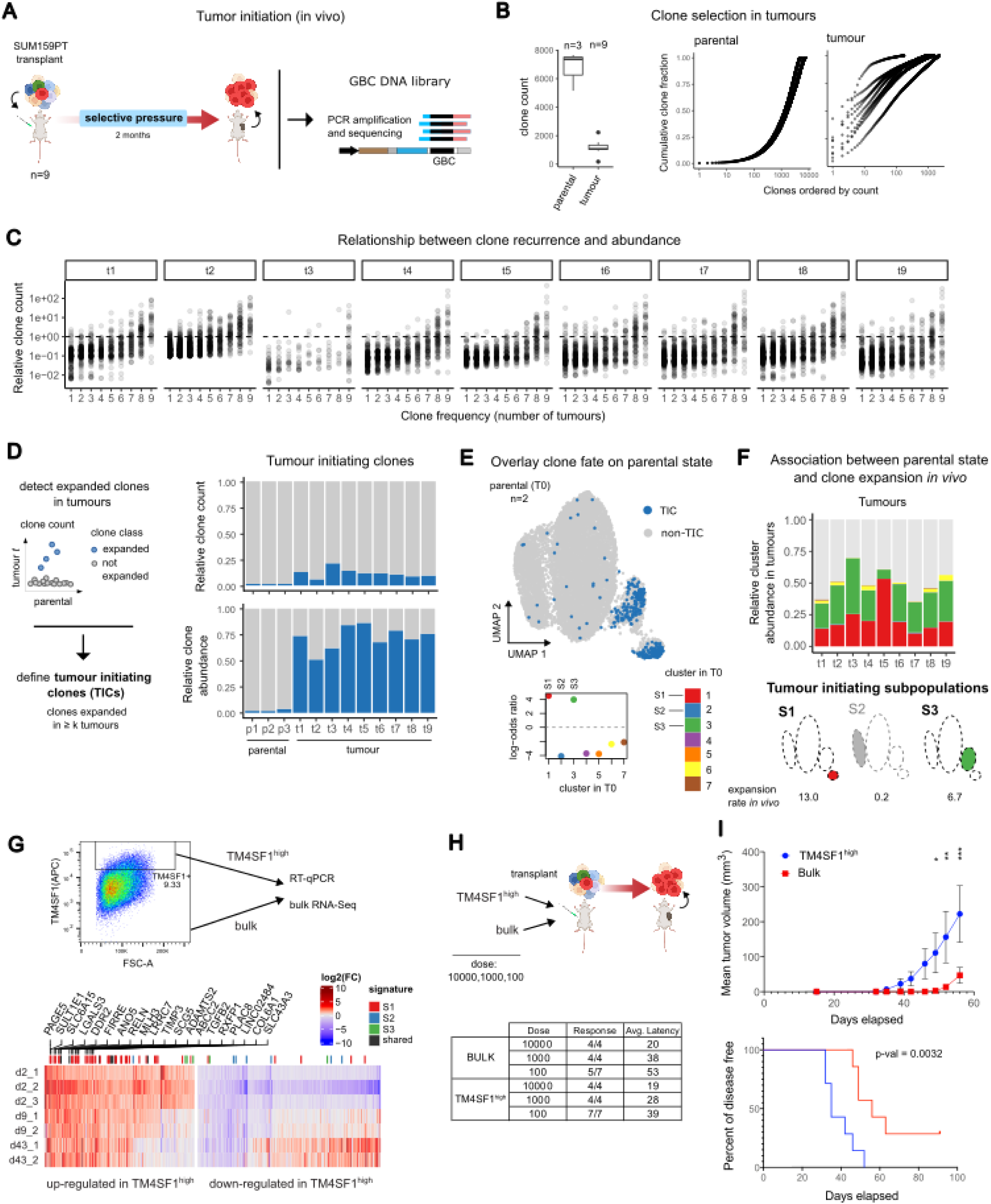
**A.** Experimental design for tumour initiation assay. Barcoded SUM159PT cells were injected orthotopically in NSG mice; after tumour formation, clonal identity and frequency were evaluated by retrieving GBC from gDNA. Three samples were also sequenced before injection (“parental”) as a control. **B.** Left: comparison of total clone count (*y* axis) between parental and tumour samples, where clones are distinct GBC species. Right: cumulative clone distribution in parental and tumour samples; for each sample, GBCs are ordered by non-increasing abundance, the *x* axis is the GBC rank, and the *y* axis is the cumulative clone frequency. **C.** Relationship between clone abundance and recurrence in the nine tumours. Each graph refers to a tumour and each dot is a clone (*i.e*., a GBC, see Methods); clones are grouped by the number of times *x* they are observed across tumours (*k*). The *y* axis is the relative clone abundance, expressed as the frequency over the total tumour size. **D.** Detection of tumour-initiating clones (TIC). Left: cartoon showing the comparison between clone abundance in a tumour *t* compared to the (average) abundance in the parental population; clones significantly more abundant in *k* = 6 out of 9 tumours are defined as tumour initiating (TIC). Right: bar plots showing the fraction of clones (top) or the relative clone abundance (bottom), in parental and tumour samples, grouped by clone class (tumour-initiating or not). **E.** Mapping of the tumour initiating clones at parental state (T0). Top: UMAP representation of T0 cells in gene expression space with cells classified as TIC in blue. Bottom: log-odds-ratio obtained from the contingency table comparing cluster assignment and TIC labelling across cells at T0. **F.** Association between parental state (T0) and clone expansion *in vivo*. Top: bar plot showing the relative normalised abundance of T0 clusters in every tumour (unassigned clones shown in grey). Bottom: cartoon highlighting TIC enrichment in subpopulations (odds-ratio values reported below). **G.** Prospective isolation of S1 cells by FAC-sorting with TM4SF1 antibody (APC-conjugated, see also Figure 1G). Top: gate used for sorting the TM4SF1^high^ population. Bottom: heatmap showing the differentially expressed genes (|log_2_(FC)| > 1 and p < 0.05) between TM4SF1^high^ and bulk cells (RNA-seq) immediately after sorting (day 0), or at 9 and 43 days from sorting of passive propagation in 2D (2 replicates each). Entries are expression log_2_(FC) between TM4SF1^high^ and bulk cells at the same time point (day 0, 9, 43 respectively). Genes are grouped into up- and down-regulated, with S1, S2, and S3 gene signatures (from Figure 1G and S7D) highlighted in colour (genes shared among multiple signatures in black). The 20 top up-regulated genes in TM4SF1^high^ cells are highlighted. **H.** TM4SF1^high^ cells are enriched for TICs. Top: tumour initiation assay using either TM4SF1^high^ or bulk cells injected orthotopically at different dilutions. Bottom: response and average latency for each dilution. **I.** Tumour growth and disease-free survival analysis in the orthotopic surgical resection model. Top: plot showing the growth dynamics (in days) of each individual primary tumour derived from transplantation of 100 cells, TM4SF1^high^ versus bulk, n=7 mice/group. Data are the mean ± SEM. P = 0.012 (49 days); P = 0.0011 (52 days); 0.000074 (56 days) by two-sided, unpaired t-test. Bottom: Kaplan-Meier curve reporting the time-dependent appearance of primary tumours derived from injection of 100 cells in mammary fat pads (Log-rank Mantel-Cox test).

### The baseline programs S1 and S3 predict tumour initiation

To determine which transcriptional states are primed to tumour initiation, we traced TICs back to their parental population. TICs were strongly associated with S1 and S3 transcriptomes at baseline, with the two subpopulations showing a similarly strong enrichment (average odds-ratio 4.4 and 4.1 for S1 and S3, respectively; **Figure 2E** and **S2C** and **Suppl. Table 4**). Both S1 and S3 were transcriptionally stable in culture in a timeframe of two weeks, as shown in **Figure 1F**, suggesting that the gene expression profile of TICs at baseline may be predictive of the phenotype. S1 and S3 showed different gene expression patterns, with S1 being clearly separated from all other clusters in gene expression space (average silhouette width 0.29 and 0.30, respectively; **Figure S1E**), suggesting that TICs stem from two distinct transcriptional programs. We then asked whether the clones in S1 and S3 differ in terms of their expansion potential *in vivo*. When the subpopulation identity was mapped on tumours, clones in S1 and S3 highly contributed to the tumour mass, relative to their initial abundance at baseline, compared to the other clones (**Figure 2F** and **S2D**); upon transplant, the expansion rate of S1 was higher compared to S3 (12.7-fold for S1 and 6.9-fold for S3, on average; **Suppl. Table 4**). We profiled the transcriptome of SUM159PT tumours by bulk RNA-Seq. Neither the S1 nor the S3 signatures as a whole were upregulated in SUM159PT tumours with respect to the parental population, including some among the top significant genes (see **Figure S2E-F** and **Suppl. Table 5**), hinting that clones undergo a transcriptional reprogramming upon transplant. This is in line with the cancer stem-cell hypothesis, where a small, stem-like, cell subpopulation exhibits both tumorigenic and differentiation potential. Of note, S100A4, one of the top significant genes in the S1 signature (see **Figure 1G**), was upregulated in SUM159PT tumours (log_2_(FC) = 2.24 and adj. p-value = 1.09e-27). In primary TNBC tumours (Pal et al. 2021) we could detect both S1 and S3 programs and identify S1^+^ and S3^+^ cell subsets accordingly (**Figure S3A-B**); in particular, we detected a strong upregulation of S100A4 in the S1^+^ subset (see **Figure S3C** and **Suppl. Table 6**). Moreover, the S100 gene family has been linked with subclonal dissemination in metastatic pancreatic ductal adenocarcinoma, with S100A4 showing the highest association (Simeonov et al. 2021) (Low et al. 2023) (**Figure S2G**). We conclude that the tumour-initiating niche of SUM159PT may share markers across different cancer diseases and, although plastic, could be partially reminiscent of its molecular state at baseline. To directly verify the tumour-initiating potential of the S1 subpopulation, we searched for surface markers for prospective isolation. TM4SF1 (transmembrane 4 L6 family member 1) was among the top 20 significant genes of the S1 signature, as well as highly upregulated in SUM159PT tumours (log_2_(FC) = 3.40 and adj. p-value = 2.85e-34, see **Suppl. Table 5**). The TM4SF1 protein regulates cell-cell adhesion and tissue invasion (Zukauskas et al. 2011); in cancer, it induces epithelial-to-mesenchymal transition (EMT), angiogenesis, migration, and self-renewal capacity (Fu et al. 2020), and its upregulation has been associated with reduced survival in TNBC (Xing et al. 2017, Chen et al. 2022). We could not identify any S3-specific surface marker. We set up a strategy for isolating TM4SF1^high^ cells by FAC-sorting (gated on top 5%, see **Figure 2G** and **Figure S4A-C**); the TM4SF1^high^ population showed extensive upregulation of several genes in the S1 signature compared to the bulk population, and this was not the case for genes in S2 and S3 signatures (RT-qPCR and RNA-seq, see **Figure 2G** and **S5A-C**). Of note, the expression of the S1 signature was maintained even after several passages in culture (see **Figure S5C and Suppl. Table 7**). TM4SF1^high^-associated genes were mainly related to invasion and metastasis pathways and suggestive of TWIST1, STAT3 and HIF1A activation (**Figure S5D**). Limiting dilution transplantation is a well-established approach to quantify the tumour-initiating content of a cell population. Therefore, we injected orthotopically serial dilutions from bulk and TM4SF1^high^ populations (**Figure 2H** and **S4A**) into NSG (NOD/SCID/IL2Rγ_c_^-/-^) immunodeficient mice. At the lowest dilution (100 cells), TIC number is a limiting factor and TM4SF1^high^ cells developed tumours with higher efficiency than mice transplanted with the same number of bulk cells (26% average latency reduction, see **Figure 2I** and **S5E-G**), suggesting that the TIC content of the TM4SF1^high^ subpopulation is higher. We conclude that S1 holds an increased tumour-initiating capacity compared to the whole SUM159PT population.

### The S3 program confers a selective growth advantage upon chemotherapy

We next investigated the response of cancer clones upon drug response *in vitro* on cultured cells and *in vivo* on transplanted tumours, using paclitaxel, an anti-mitotic chemotherapy agent used to treat many cancer types (Weaver 2014). We treated barcode-labelled SUM159PT cells at 50 nM (which corresponds to ∼IC95, **Figure S6A**) for three days in culture, with the untreated condition as a control (see **Figure 3A**, top). To evaluate the drug response *in vivo*, we transplanted barcode-labelled SUM159PT cells into the mammary fat pads of six NSG immunodeficient mice; once tumour was formed, mice were treated with paclitaxel every 5 days (see **Figure 3A**, bottom). *In vitro*, treatment induced a deep clonal selection: between 9% and 22% of the initial clone pool survived ≥ 10 days post paclitaxel removal, while cells cultured for a comparable time span in the absence of treatment did not undergo clonal selection (see **Figure 3B**, top, and **S6B** and **Suppl. Table 4**). A comparable effect was observed on an independent barcoding experiment (see **Figure S6C**). *In vivo*, paclitaxel treatment delayed tumour growth, but did not trigger remission of the disease (see **Figure S6D**); the major driver of clone selection was the tumour-initiation capacity (see **Figure 3B**, bottom). Of note, the highly expanded surviving clones were not randomly selected by treatment, both *in vitro* and *in vivo* (see **Figure S6E-F)**; therefore, we reasoned that both survival and proliferation potential upon paclitaxel treatment were pre-encoded. We defined the *drug-tolerant clone* (DTC) pool as the set of clones that were significantly more abundant after treatment in at least 4 out of 6 samples compared to their average abundance at basal state (Fisher’s exact test, see Methods and **Figure 3C**). We detected 171 and 164 DTCs *in vitro* and *in vivo*, respectively. When traced back to their baseline transcriptional state, clones surviving drug insult *in vitro* were depleted in S1, but were more abundant in S3 than expected by chance (see **Figure 3D**, left and **S7A,C** and **Suppl. Table 1**). In contrast, clones surviving drug treatment *in vivo* were belonging to either S1 or S3 (see **Figure 3D**, right and **S7B-E** and **Suppl. Table 1**). Note that 71% of TICs were also drug-tolerant *in vivo* (see **Figure S7D-E** and **Suppl. Table 4**), confirming that the effect of paclitaxel *in vivo* was modest. When assessing the relative abundance of S1 and S3 clones in tumours treated with paclitaxel, S3 showed a higher fitness over S1 (**Figure 3E** and **S7E** and **Suppl. Table 4**), in agreement with *in vitro* results. We deduced that the drug tolerance phenotype is different from the tumour-initiating capacity in SUM159PT and is primarily associated with the S3 baseline program.

**Figure 3.**
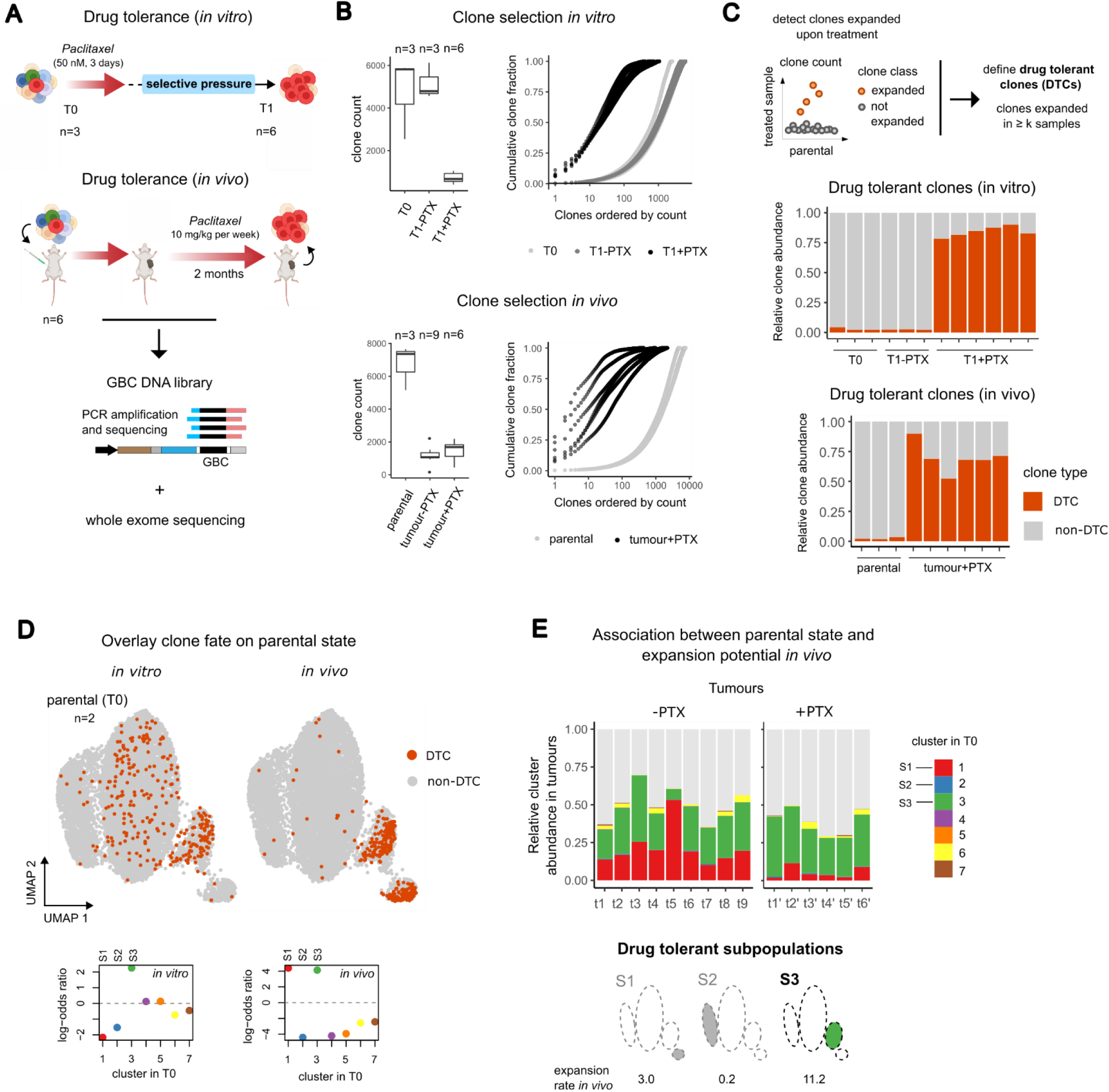
**A.** Experimental design, drug tolerance assay. Top: *in vitro* assay. Barcoded SUM159PT cells were treated with paclitaxel *in vitro* and harvested when single cell colonies were grown (n=6). GBC loci were PCR amplified and sequenced. The untreated parental population at T0 (n=3) and T1 (n=3) was also sequenced as a control. Bottom: *in vivo* assay. Barcoded SUM159PT cells were injected orthotopically NSG mice; after tumour formation, mice were treated or not (see Figure 2A) with paclitaxel. Parental samples (n=3) were also sequenced as a control. **B.** Clone selection upon treatment. Top: comparison of total clone count (left) and cumulative clone distribution (right) in parental, untreated, and treated *in vitro* samples. Bottom: same as above, for parental, untreated tumour (as in Figure 2A), and treated tumour samples (see also Figure 2B legend). **C.** Detection of drug tolerant clones (DTC). Top: cartoon showing the comparison between clone abundance in each sample compared to the (average) abundance in the parental population; clones significantly more abundant in *k* = 4 out of 6 samples are defined as drug tolerant. Bottom: bar plot showing the relative clone abundance in parental and treated samples *in vitro* and *in vivo*, respectively, and grouped by class (drug tolerant or not). **D.** Mapping of the drug tolerant clones at parental state (T0). Top: UMAP representation of T0 cells on gene expression space, with cells classified as DTC *in vitro* (left) or *in vivo* (right) coloured in orange. Bottom: log-odds-ratio obtained from the contingency table comparing cluster assignment and DTC labelling *in vitro* (left) or *in vivo* (right) across cells at T0. **E.** Association between parental state (T0) and clone expansion *in vivo*, without and with treatment. Top: bar plot showing the relative normalised abundance of T0 clusters in every untreated (left, as reported in Figure 2F) or treated tumour (right), respectively (unassigned clones shown in grey). Bottom: cartoon highlighting the subpopulations enriched in DTCs (odds-ratio values reported below).

### GALILEO links cancer clones with transcriptional programs and DNA accessibility states at single-cell resolution

To investigate the epigenetic state of cancer clones and relate it to the transcriptional readout, we developed GALILEO, Genomic bArcoding pLus sIngLE-cell multi-Omics. Specifically, we performed single-cell multiome ATAC plus gene expression sequencing on barcode-labelled SUM159PT nuclei in two biological replicates (**Figure 4A**). This assay enables access to gene expression, DNA accessibility, and clone information simultaneously at single-nucleus resolution. At baseline, we identified 2023 to 2024 unique clones and assigned a clonal identity to between 73% and 86% of high-quality nuclei (see **Figure S8A** and **Suppl. Table 1**). We obtained 7 and 6 clusters for the two replicates, respectively, and retrieved the subpopulations S1, S2, and S3 previously identified by scRNA-Seq (see **Figure 4B** and **S8B-C**), with largely overlapping genes (see **Figure S8D** and **Suppl. Table 8**). When evaluating their DNA accessibility state, nuclei from S1, S2, and S3 showed distinct profiles in the scATAC-seq space, comparable to the scRNA-seq results (see **Figure S8E**). To make sense of the >10^5^ accessible regions detected from scATAC-Seq and identify the relationships among them, we first determined a small number of *topics* [see (Bravo Gonzalez-Blas et al. 2019)], each assigning a probability value to all ATAC regions in the dataset. Next, we compared the region probability of every topic pair across the two replicates using the irreproducible discovery rate (IDR, see Methods and **Figure S9A**). We defined a subset of reproducible regions (*i.e.*, satisfying IDR < 0.05) for each topic pair, referred to as *ATAC module* hereafter, and assigned a *reproducibility score* to them (see **Figure 4C**, left and **S9B** and **Suppl. Table 9**). Note that our approach based on topic modelling discards ubiquitously accessible regions; combined with IDR filtering, this results in a substantial reduction of the size of the dataset (see pie chart on **Figure 4C**). Most reproducible regions were found in few, large ATAC modules containing more than 400 regions, the largest one containing 1511 regions (see **Figure 4D** and **S9C**). This few-to-few mapping across replicates suggests that the grouping of the regions into ATAC modules is not random. Therefore, each of these modules is expected to identify a pool of genomic elements that jointly participate in the regulation of gene expression. More than 90% of the regions could be assigned a regulatory element, according to the ENCODE cCRE registry: 9% were annotated as promoter-like signatures (PLS), within 200 bp of transcription start sites (TSS) of genes, 7% and 75% as proximal and distal enhancer-like signatures (pELS, dELS), respectively (**Figure 4C**).

**Figure 4.**
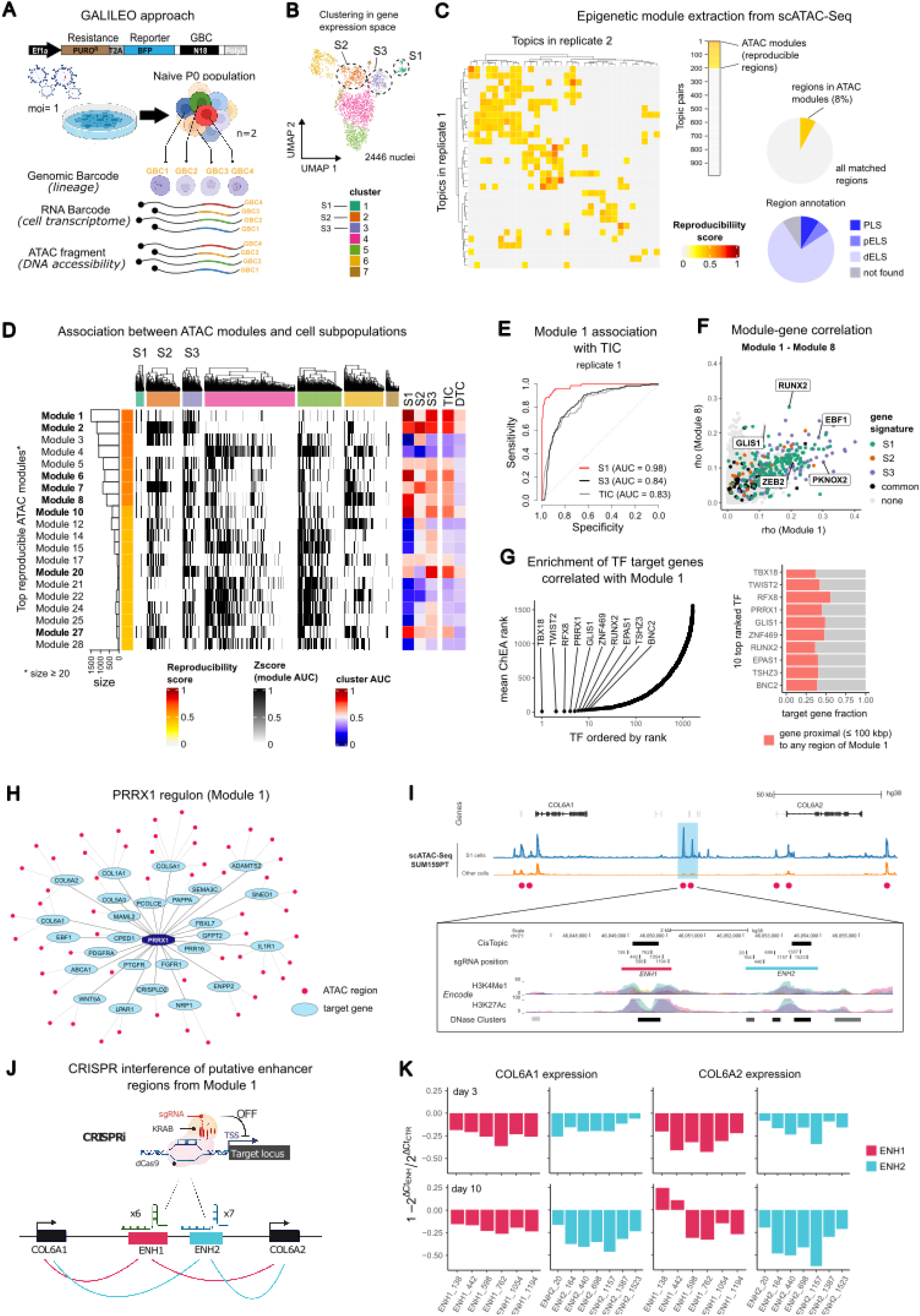
**A.** Genomic bArcoding pLus sIngLE-cell multi-Omics (GALILEO) strategy. SUM159PT infected with the GBC library are processed with the Single Cell Multiome ATAC + Gene Expression kit. The readout of each cell is it lineage, captured by genetic barcodes (GBCs), gene expression profile, captured by 3’ end RNA sequencing, and DNA accessibility state, captured by ATAC. **B.** UMAP representation of replicate 1 n gene expression space coloured by cluster (2446 nuclei); the three subpopulations are marked with dashed circles. **C.** Topic modelling on ATAC-Seq regions. Left: heatmap showing the comparison of the output of topic modelling on the two Multiome replicates; rows are topics in replicate 1, columns are topics in replicate 2, and entries are reproducibility scores (see Methods); topics displaying strong correlation with read coverage are not shown (see **Figure S8B**); topic pairs that do not share any reproducible region are coloured in grey. Centre: entries of the matrix ordered by non-increasing reproducibility score; the yellow fraction represents the ATAC modules used hereafter. Right: pie charts showing the fraction of reproducible regions in replicate 1 (top) and their annotation via the ENCODE registry (bottom) of candidate cis-Regulatory Elements (cCREs) as either PLS (promoter), pELS (proximal enhancer), and dELS (distal enhancer). **D.** Comparison between ATAC modules and gene expression in single nuclei (replicate 1). In the heatmap, rows are the top 20 scoring modules, annotated with their size (bar plots on the left), columns are nuclei split by cluster, and entries are module AUC scores representing the overall accessibility of a module in each cell (see Methods). The degree of association (AUC) of a module to subpopulations (S1, S2, and S3) and cancer fates (TIC and DTC *in vitro*), ranging from negative (blue) to positive associations is reported on the right (red). ATAC modules predicting either S1, S2, or S3 with AUC > 0.75 are reported in bold. Hierarchical clustering of columns is performed with `complete` method from `hclust` on Euclidean distances. **E.** ROC curves showing the performance of Module 1 AUC as a predictor of S1 (red), S3 (black), or TICs *in vitro* (grey) on replicate 1. **F.** Association between module AUC scores and gene expression (replicate 1). Each dot is a gene and its value represents the (positive) Spearman’s rho correlation coefficient between its expression and module AUC score, for Module 1 (*x* axis) and Module 8 (*y* axis). Genes are coloured according to whether they belong to either S1, S2, or S3 gene signatures (scRNA-Seq or multiome). Transcription factors with rho ≥ 0.2 in either module are labelled. **G.** Transcription factor enrichment on the genes whose expression is positively correlated with Module 1 AUC. Left: the *x* axis reports the TFs sorted by non-decreasing rank, the *y* axis reports the rank, and the top 10 ranked TFs are labelled. Right: fraction of genes (coloured in pink) whose locus is located ≤ 100 kbp away from any region in Module 1, for the 10 top-ranked TFs. **H.** The “PRRX1 regulon” includes the set of genes whose expression is positively correlated with Module 1 AUC and that i) are predicted as PRRX1 targets by ChEA3 and ii) lie at ≤ 100 kbp from any region in Module 1. **I.** COL6A1 and COL6A2 loci; shown are the scATAC-Seq peaks (aggregate signal, replicate 1). Red dots label Module1-specific regions. In the magnification is shown the region containing the two enhancers, ENH1 and ENH2, together with ENCODE regulatory tracks for H3K4Me1, H3K27Ac and DNase clusters and the position of sgRNAs for CRISPRi (see also **K**). **J.** Scheme showing the CRISPRi approach and the model of Enhancer-Gene pair regulation. **K.** RT-qPCR data showing the impact in COL6A1 and COL6A2 expression in TM4SF1^high^ SUM159PT_KRAB cells observed upon expressing sgRNAs targeting ENH1 (in pink) or ENH2 (in blue) (see also I). Data are measured at 3 days (top) or 10 days (bottom) after dCas9-KRAB induction by doxycycline (n=6 and n=7 sgRNAs, respectively) and compared to a group of non-targeting sgRNAs (n=4).

### ATAC modules recapitulate the multiple DNA accessibility profiles of gene expression clusters

The regions found in the same ATAC module are accessible in the same cells, and, by definition, are reproducible across replicates. We investigated the relationship between ATAC modules and transcriptional or clonal subpopulations. We computed a score for each ATAC module *M* and for each cell *c* (AUC, see Methods), representing the overall accessibility of the regions of *M* in *c,* and referred to as *module AUC* hereafter. Using module AUC, we associated several large ATAC modules with either S1, S2, or S3 (see **Figure 4D** and **S9C**). Among the 20 highly reproducible modules, 8 (highlighted in bold in **Figure 4D**) could be associated to any of S1, S2, or S3, with Module 1 (1511 regions) being the top predictor of both S1 (AUC = 0.98) and S3 (AUC = 0.84) (**Figure 4E**). Of note, the small Module 20 (50 regions) showed an equally strong association with S3 (AUC = 0.84). To further assess the relationship between ATAC modules and subpopulations, we correlated genome accessibility (using module AUC, see above) with gene expression across cells (Spearman’s *ρ*, see **Figure 4F and Suppl. Table 10**). This approach ensures that mechanisms involving either *cis* or *trans* genomic elements can be captured, as no constraint on region-gene proximity was used. Overall, module AUC was positively correlated with the expression of genes of the associated subpopulation signature (see **Figure 4F** and **S9D**). These results support the hypothesis that ATAC modules contain regulatory elements jointly involved in the control of specific transcriptional programs.

### Tumour-initiating clones share a common chromatin priming state

We then sought to investigate the role of ATAC modules in cancer phenotypes, namely, tumour-initiation capacity and drug-tolerance. Importantly, Module 1, the top reproducible accessibility state, predicted TICs with high specificity and sensitivity (see **Figure 4E** and **S9E**), consistently with it being associated with S1 and S3, which, in turn, are enriched in TICs (see also **Figure 2E-F**). This suggests that the tumour-initiating capacity may be linked to a specific and pre-existing epigenetic state and may explain the phenotypic relationship between the cells of S1 and S3. Subsequently, we used transcriptional and epigenetic information jointly and at single-cell resolution to highlight the gene regulatory networks and the epigenetic determinants involved in the tumour-initiating capacity. For each module *m*, we used the set *G_m_* of positively correlated genes (see above) as input to measure the transcription factor activity in *m* given a set of putative targets for each TF (Lachmann et al. 2010) (**Figure 4G**, **S9F and Suppl. Table 11**). Among the top ranked TFs for Module 1, we detected several TFs that have been previously linked to tumour-initiation capacity, including TWIST2, PRRX1 and RUNX2. TWIST2 is a member of the TWIST family of TFs, which has been extensively associated with poor tumour prognosis, epithelial-to-mesenchymal transition (EMT), and stem-cell activity in breast cancer (Ansieau et al. 2010, Beck et al. 2015, Nobre et al. 2022). Similarly, RUNX2 activity has also been linked to EMT and to invasive phenotypes (LaFave et al. 2020), which lead to metastasis, notably in the breast (Watson et al. 2021, Brown et al. 2022); finally, PRRX1 has been recently shown to sustain metastatic dissemination and induce a switch to a mesenchymal-like state in a melanoma cancer model (Karras et al. 2022). Of note, TWIST2 and PRRX1 resulted as top ranked predicted in Module 1 also using the set of genes proximal to Module 1 ATAC regions as input (**Figure S9F**). For each TF, we identified its regulon by the set of positively correlated target genes whose locus is proximal (< 100 kbp) to any region of Module 1. Of note, several genes in the S1 signature, including Procollagen C-endopeptidase enhancer 1 (PCOLCE) and collagen-encoding genes (COL6A1, COL6A2, COL5A1, COL5A3), were found as part of PRRX1 and TWIST1 regulons (**Figure 4H and S9G)**. To directly verify the role of the genomic elements of Module 1 in gene regulation, we selected two regions located in the proximity of COL6A1 and COL6A2, highly accessible in S1 cells (**Figure S9H**) and classified as dELS by ENCODE cCRE (**Figure 4I**, top). Subsequently, we targeted the two regions by means of an inducible CRISPR interference strategy (Gilbert et al. 2013, Larson et al. 2013) (**Figure 4J**). We observed that repression of either region led to a consistent and reproducible reduction in COL6A1 (up to 36% and 46%) and COL6A2 expression (up to 43% and 62%; **Figure 4K**), both at early (3 days) and late time points (10 days) post-dCas9-KRAB induction.

### A subset of the drug tolerant subpopulation exhibits a pre-existing genomic amplification

Module 20 predicts S3 with high specificity and sensitivity (AUC=0.84 and 0.75 in the two replicates, see **Figure 5A** and **S10A**). As shown in **Figure 3**, the S3 program is associated with increased drug tolerance, both *in vitro* and *in vivo*. We noticed that most regions of Module 20, as well as several genes of the S3 signature, were located on a 5.5 Mbp-long region of chromosome 11 (**Figure S9I**). The odds are low that an epigenetic regulatory mechanism involves a cluster of highly localised genomic elements and thus we reasoned that a genetic alteration might better explain the transcriptional profile of S3. When interrogating the whole-exome sequencing profile of paclitaxel-treated samples and comparing it to that of untreated cells (n = 3, see **Figure 3A**), we detected 18 recurrent copy number variants (CNVs, see **Figure 5B** and **S10B**). Notably, the top amplified region (average log_2_(FC) = 0.53 and p-value = 5.65e-185) lied on chromosome 11, specifically across bands 11q23-11q24 (**Figure 5C**). These results suggest that the amplification was already present in S3 cells before treatment and that Module 20 captures a specific genetic background of S3, rather than a localised increase in chromatin accessibility. This implies that the drug tolerance phenotype is, at least in part, genetically determined, and, in turn, suggests that a subset of DTCs could maintain a stable memory of the treatment. Therefore, we investigated the susceptibility of cells to paclitaxel upon a recovery period of 24 days after a first round of treatment (see Methods, **Figure 5D**, top, and **S10C**). Upon a second round of treatment, clonality was reduced by 63% (average recovery at day ≥ 17 compared to day ≤ 3, see **Figure 5D**, bottom), suggesting that chemotherapy was still effective; however, the drug sensitivity resulted lower compared to a single round of treatment, where only 19% of clones survived. Finally, to verify the specificity of the association observed between the chr11 region amplification and resistance to paclitaxel, we examined a drug resistance model, where SUM159PT cells were repeatedly treated with increasing doses of paclitaxel (see Methods, **Figure 5E**, top, and **Figure S10C**) up to the onset of a drug resistant phenotype. The WES profile confirmed an amplification on chromosome 11 whose locus overlap 75% of the above detected one (**Figure 5E**, bottom, **S10D** and **Suppl. Table 12**). We conclude that, in SUM159PT, paclitaxel-based chemotherapy causes a clonal expansion of a subpopulation harbouring the amplification of 11q23-11q24.

**Figure 5.**
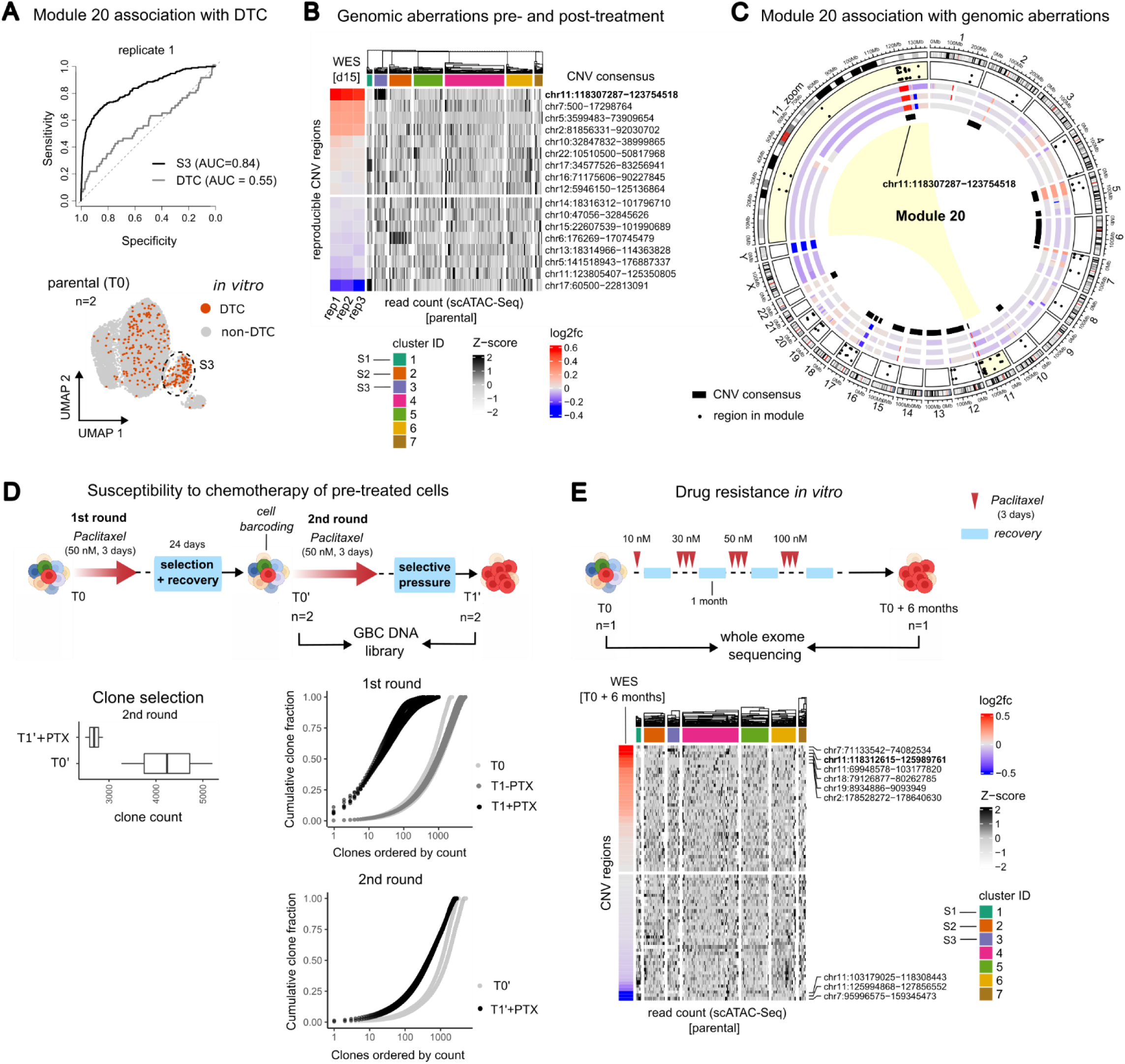
**A.** ROC curves showing the performance of Module 20 AUC as a predictor of S3 (black) or DTCs *in vitro* (grey) on multiome replicate 1. Bottom: UMAP representation of T0 cells on gene expression space coloured as in Figure 3D. **B.** Copy-number variants (CNVs) pre- and post-treatment (drug tolerance assay, replicate 1). Heatmap showing the association between ATAC-Seq signal from multiome data and copy-number variants inferred by WES. Rows are consensus CNV obtained from the analysis of paclitaxel-treated samples (day 15; n=3 replicates), as in the in vitro experimental design shown in Figure 3A; columns are nuclei in multiome replicate 1; entries are cumulative ATAC counts in each CNV locus and in each nucleus. Rows correspond to consensus CNVs across three replicates. The coverage log_2_(FC) between each treated sample and the untreated reference (baseline SUM159PT cells) is reported on the left; chromosome location is shown on the right. Columns are grouped by gene expression cluster; hierarchical clustering is performed with `complete` method from `hclust` on Euclidean distances. **C.** Circos plot showing the association between regions of Module 20 (see Figure 4C) at baseline and CNVs in treated condition. In the dotplot (external ring), the x axis is the location on the genome and the y axis is IDR for the regions in Module 20; all CNVs for the 3 replicates are shown and coloured according to the log_2_(FC) (middle ring); consensus CNVs are marked in black (internal ring). **D.** Drug tolerance assay second round. Top: SUM159PT cells were treated with paclitaxel and clone selection was stabilised until T0’; cells were subsequently infected with the library of Genetic Barcodes (GBC), sorted for BFP expression, subjected to a second round of treatment, and harvested at T1’ when single cell colonies were grown. The three populations (T0, T0’ and T1’) were sequenced. Bottom left: comparison of the total clone count at T0’ and T1’. Bottom right: cumulative clone distribution in untreated and treated samples after either 1 round (see also Figure 3B, top) or 2 rounds of treatment. **E.** Top: Experimental design, long-term drug resistance assay *in vitro*. SUM159PT cells were repeatedly treated with increasing doses of paclitaxel as shown until resistant clones where obtained. The exome of untreated and resistant clones was then sequenced. Bottom: detection of genomic aberrations pre- and post-treatment (replicate 1). The heatmap shows the association between the ATAC-Seq signal from multiome data and copy-number variants predicted by WES (see also **B**); rows are the CNVs obtained from the analysis of paclitaxel-treated samples (n=1) and are annotated with the log2(FC); entries are the cumulative ATAC counts in each CNV locus, as in **B**.

### Two distinct clone lineages can enter a drug tolerance state

We note that the 11q23-11q24 amplification is only found in S3. However, *in vitro*, many DTC clones fall outside S3 (see **Figure 5A** and **S10A**), highlighting their molecular heterogeneity at the baseline. To further characterise the pathways leading to drug tolerance *in vitro*, we designed a single-cell time-course experiment on barcoded SUM159PT cells (**Figure 6A**, top). To minimise technical variability, we treated cells for 3 days with Paclitaxel, collected samples every two days after drug removal (day 5 to 15) and sequenced them simultaneously using a reverse time course as in (Loukas et al. 2023). Consistent with untreated samples, we assigned a clonal identity to most of the high-quality cells (80-84%, see **Figure S11A-B**). Treatment elicited a progressive clone selection, from 1126 distinct clones at day 5 to 433 at day 15 (**Figure 6A** and **Suppl. Table 13**). An independent experiment confirmed the clone selection dynamics (**Figure S11B-C**). Chemotherapy induced a substantial transcriptional change (**Figure 6B-C** and **S11C-D** and **Suppl. Table 14**). At initial time points (d5-d9), most cells were drug-sensitive and developed two distinct responses to treatment: the one characterised by the induction of stress response pathways, including amino-acid deprivation, unfolded protein response and inflammation (cluster 1); the other mostly characterised by autophagy (cluster 2, **Figure 6D** and **S11E** and **Suppl. Table 15**). Conversely, at late time points cells surviving the treatment showed enhanced translation activity (cluster 3). At initial time points (d5 and d7), cells belonging to the surviving pool (*i.e.*, DTCs) accounted for only 9% of the whole sample and their transcriptional profile was scattered (**Figure 6B** and **S11D)**. DTCs survived the treatment by remaining in a state of suspended proliferation for several days and, starting from days 9-11, entered an intense proliferation phase; from day 13 on, DTCs constituted 73% to 85% of surviving cells (**Figure 6A** and **S11C**) and acquired a distinguishing transcriptional profile (**Figure 6E**). At late time points (days 13-15), cells stemming from the same clone at baseline overall displayed a divergent transcriptional program (solid line), comparable to that of cells belonging to different clones (dashed line, **Figure 6F**). Then, we asked whether any distinguishing transcriptional footprint exists in highly expanded clones. To do this, we devise an unsupervised approach to find sets of mutually similar clones (by defining a *pair propensity* score across gene expression neighbourhoods, see Methods and **Figure 6G**). Two groups of clones, or *lineages*, showed mutually high transcriptional similarity (**Figure 6G-H** and **S12A**), suggesting that multiple pathways to drug tolerance may exist in SUM159PT cells. Both lineages contained highly abundant clones, indicating that the transcriptional readout is not associated with a higher or lower proliferation potential. The two lineages were reproducible both across time points and across independent experiments (**Figure S12B-C**). On average, they accounted for 50% (lineage 1) and 35% (lineage 2) of the cells at late time points (the remainder fraction belongs to unclassified clones). Notably, clones in lineage 1 stemmed from S3 and were characterised by a pre-existing genomic amplification on band 11q23-11q24. Among the upregulated genes in lineage 1, we detected MT1E, whose relevance in breast and other cancer types has been extensively proven (Masiulionyte et al. 2019, Wang et al. 2021), as well as FEZ1 and RPS25, top significant genes in the S3 signature (**Figure 6I-J** and **Suppl. Table 14**). Consistently, genes located within the 11q23-11q24 amplification, which is specific to S3, were upregulated in clones belonging to lineage 1 (**Figure 6K** and **S12D-E**). This showed that the transcriptional differentiation breadth of the two lineages is determined by the genetic background of the ancestor clone, specifically, depending on whether it carries the 11q23-11q24 amplification or not. In contrast, we did not detect any lineage 2-specific copy-number aberration (**Figure S12F**). The top upregulated genes in the lineage 2 signature, namely, S100A2, IGFBP2, IFI27, and PVT1, were most highly expressed immediately following treatment in both DTCs and non-DTCs (**Figure 6J**), suggesting that the transcriptional program of lineage 2 is not DTC-specific. Of note, the early response to paclitaxel includes upregulation of PVT1, a long non-coding gene acting as negative regulator of the transcription factor MYC, a key regulator of growth and cellular metabolism, frequently associated to breast cancer (Cho et al. 2018). Consistently, we observed that MYC activity decreased immediately after treatment and increased during adaptation in response to it (**Figure S12G**), with only slight differences between the two lineages (**Figure S12H**). Of note, and consistent with our findings, recent evidence showed that a reduced MYC activity promotes a chemotherapy survival phenotype in breast cancer via the adoption of an embryonic-like diapause state (Dhimolea et al. 2021). In conclusion, we mapped at clonal level the transcriptional response upon drug treatment and isolated different pathways of cancer transcriptional evolution leading to resistance, one being invariably linked to a pre-existing genetic rearrangement.

**Figure 6.**
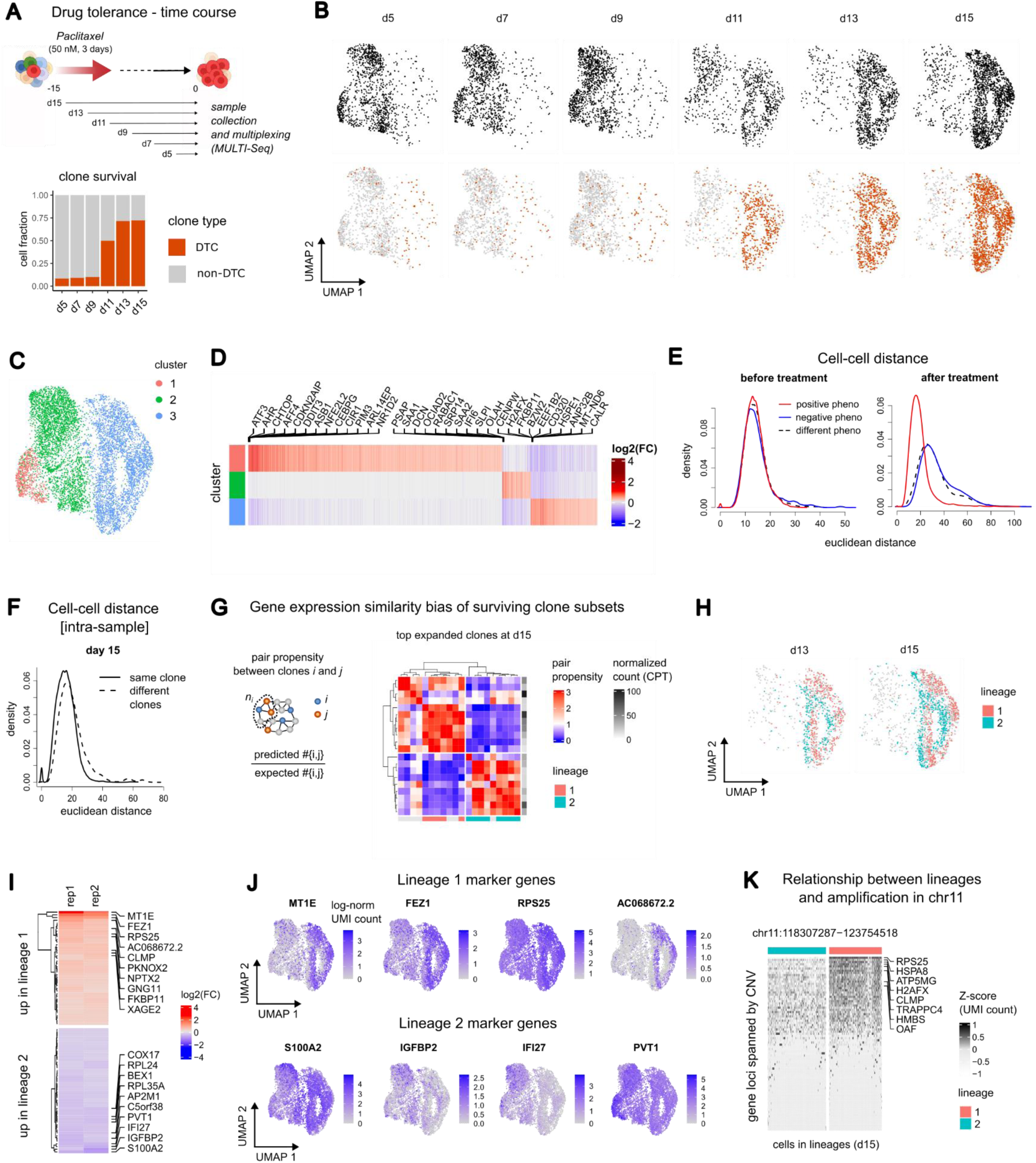
**A.** Experimental design, time-course drug tolerance assay (replicate 1). Top: SUM159PT cells were treated with paclitaxel for 3 days; cells were harvested every 2 days post-paclitaxel removal (n=1 per time point) and processed with scRNA-Seq using a reverse time-course multiplexing strategy (MULTI-Seq, see Methods). Bottom: bar plot showing the distribution of DTC *in vitro* (as defined in Methods and shown in Figure 3C, top) on treated samples. **B.** Drug-tolerant clone selection across time. Cells from treated samples are mapped to a common gene expression UMAP space (7884 cells in total) and split by sample (top) or coloured according to whether they are drug tolerant (in orange) or not (in grey). **C.** UMAP representation of cells at day 5, 7, 9, 11, 13, and 15 (exp1) and coloured according to cluster assignment. **D.** Cluster signatures (exp1). Heatmap where rows are genes, columns are clusters, and entries are log_2_(FC) values between a cluster and its complement. Columns are annotated with the same colour as in **C**. Significantly up-regulated genes in any of the three clusters are shown and order by non-decreasing average rank between the log2(FC) (higher to lower) and the p-value (lower to higher). The top 10 ranked genes for each cluster are labelled. **E.** Gaussian kernel density of Euclidean distances between DTCs (red line, positive phenotype), non-DTC (blue line, negative phenotype), and between DTCs and non-DTCs (dashed black line, different phenotype) computed on gene expression space either before (T1, left) or after treatment (exp1, right). **F.** Gaussian kernel density of Euclidean distances between sister cells (solid line) and non-sister cells (dashed line) computed on day 15 sample (see Methods). **G.** Evaluation of gene expression similarity bias by pair propensity. Left: schema illustrating the calculation of the pair propensity measure between clones (see Methods). Right: pair propensity value for top expanded clone pairs at day 15 (clones *i* with *p_ii_* < 1 are not shown); rows and columns are distinct clones, rows are annotated by clone abundance (in CPT, count per thousand cells) and columns are annotated by lineage. **H.** UMAP representation of cells at day 13 and 15 coloured according to lineage assignment (lineage 1 in pink, lineage 2 in cyan, and unassigned clones in grey). **I.** Heatmap where rows are lineage gene signatures, columns are replicates, and entries are log_2_(FC) values between lineage 1 and lineage 2. Genes are ordered by non-increasing log_2_(FC) in replicate 1. The 10 DEGs with respectively higher and lower log_2_(FC) are labelled. **J.** Lineage signatures (replicate 1). UMAP plot of cells at day ≥ 5 coloured by log-normalised gene expression of the 4 top log_2_(FC) genes of lineage 1 (top) and lineage 2 (bottom). **K.** Heatmap where rows are genes whose locus is spanned by the top amplified CNV in drug treated samples (see Figure 5A), columns are cells at day 15 and entries are log-normalised, scaled UMI counts. Rows are sorted by non-increasing average expression across all cells. Columns are split by lineage and clustered with `complete` method on Euclidean distances.

## DISCUSSION

One of the main challenges in cancer biology is predicting how tumours evolve in response to changes in the tumour environment. The capacity of one or more clones to sustain tumour growth at distal sites or to trigger disease relapse upon cytotoxic treatment may depend on a specific set of molecular characteristics. Their identification has been the focus of intense research efforts, both for the obvious clinical applications and for the understanding of the mechanisms underlying tumour plasticity.

Recently, it has been suggested that the cancer stem-like pool might be heterogeneous, with distinct subpopulations primed to different fates (Lapidot et al. 1994, Bonnet and Dick 1997, Al-Hajj et al. 2003, Singh et al. 2004, Ricci-Vitiani et al. 2007, Lathia et al. 2015, Batlle and Clevers 2017). The SUM159PT model is one representative example, in which distinct states coexist in equilibrium in the same cancer cell population (Gupta et al. 2011). Here, we provided the distinctive transcriptional and epigenetic traits of each sub-pool. To the best of our knowledge, this is the first time that tumour initiation and drug tolerance were analysed at single-cell level on lineage-barcoded cells and on the same system, providing a high-resolution representation of tumour complexity.

We initially asked which clonal subpopulations lie in a defined transcriptional or epigenetic state. In the case of SUM159PT, only a fraction of the clones displayed stable transcriptional profiles at the subpopulation level, hinting that these may encode for specific functions. Indeed, the tumour initiation clones (TICs) were almost exclusively associated with either S1 or S3 signatures, with S1 showing the strongest enrichment and S3 also encoding for drug tolerant clones (DTCs). The third signature (S2) was not attributable to any cancer property, although it contains basal markers (*e.g*., miR-205HG, HMGA1).

An important and direct conclusion of our study is that TICs and DTCs do not coincide but coexist in the same cancer population, sharing a minor subset of clones (belonging to S3). TICs and DTCs can significantly change their transcriptional profile during cancer evolution, adapting to the environment and conditions. In transplanted tumours, TICs lose their baseline signature upon expansion and maintain upregulation of only a handful of markers (*e.g*., S100A4, TM4SF1), whose relevance in the phenotypes is confirmed in the literature (Simeonov et al. 2021, Hou et al. 2022, Low et al. 2023). Similarly, DTCs undergo a massive transcriptional reprogramming after treatment and, thus, are strikingly distinct from their non-DTC counterpart, a behavior which is similar to other cancer cell models (Dhimolea et al. 2021, Loukas et al. 2023). Using lineage tracing information on a time-course single-cell profiling, we could describe two distinct and co-occurring transcriptional trajectories in drug adaptation. Differently from TICs, the DTC subpopulation did not show a strong transcriptional or epigenetic determinant at the baseline. However, a subset of DTCs, lying in S3, shows amplification of a 5.5 Mbp-long region of chromosome 11 (bands 11q23-11q24). This genetic background also reproducibly segregated with chronic treatment resistance in the SUM159PT model. To our knowledge, this amplification is not reported as a recurrent alteration in cancer (https://cancer.sanger.ac.uk/cosmic) and the relevance of the individual genes contained within the locus remains to be evaluated. However, the 11q23-11q24 locus contains the miRNA cluster MIR100HG, which encode for miR-125b, a miRNA known to confer resistance to taxol treatment in TNBC cell lines (Zhou et al. 2010).

A key innovative element of our study is represented by the use of a cutting-edge approach that combines single-cell multi-omic profiling (transcriptome and DNA accessibility) with lineage tracing (clone information at an arbitrary time *t*, which we call the baseline – P0). Specifically, we found a putative regulatory program (Module M1) common to both S1 and S3, which elegantly links the two tumour-initiating states. Moreover, the inferred transcription factors and the corresponding regulons comprise both genes and regulatory regions, including non-coding components (lncRNAs and enhancers). Note that these elements might be detectable in bulk experiments, but only the single-cell resolution can explain their relationship, which may be subpopulation-dependent. The TF hubs of the predicted regulons are fully supported by the literature; for instance, PRRX1, TWIST2 and RUNX2 have been linked to breast cancer and to the epithelial-to-mesenchymal transition (EMT), a founding element shared by both tumour aggressiveness and stem cell identity programs (Dave et al. 2012, Roche 2018).

An open question is whether the molecular traits distinctive of TIC are specific to SUM159PT or are generalisable. By employing either lineage tracing or single-cell omics, recent literature highlighted programs with remarkable similarities to the ones we reported. In a genetically engineered mouse lung cancer model, a co-accessibility module characterized by RUNX2 activity was identified and linked with the acquisition of a pre-metastatic state, and the associated genes were predictive of the outcome of human lung cancer (LaFave et al. 2020). Accordingly, single-cell lineage-tracing revealed the underlying program of a pool of metastatic initiating cells in melanoma, characterised by high PRRX1 expression and promoting the establishment of a mesenchymal-like cell state (Karras et al. 2022). Finally, the KP-tracer approach for *in vivo* lung cancer lineage tracing allowed to show that tumours evolve through stereotypical trajectories, with the transient activation of cellular plasticity programs and a subsequent clonal sweep of highly fit sub-populations marked by an early or late mesenchymal transition (Yang et al. 2022).

Our and published evidence suggests a model where the mechanisms influencing clonal fate and tumour evolution tend to converge towards a common epigenetic state, often established before a challenge and, therefore, predictable. We foresee that combining cutting-edge molecular tools at the genome scale, like the ones presented here, as well as genetic ones, with suitable computational frameworks, could critically contribute to further dissect the role played by different transcriptional, epigenetic and genetic layers in cancer evolution. Our study has made it clear that a multi-layered framework is feasible and an invaluable resource to this end.

## ACKNOWLEDGEMENTS

We thank Chiara Tordonato for help with mice experiments; Leah Rosen and Magdalena Strauss for input on barcode analysis; Pier Giuseppe Pelicci, Niccolò Roda and Valentina Gambino for help with the Perturb-seq lentiviral infection. We acknowledge support by the technological units at the European Institute of Oncology (IEO), in particular to the Genomic Units and Luca Rotta, the Sorting Service and Simona Ronzoni, the tissue culture facility, the imaging unit and the bioinformatics unit; the EMBL-EBI gene expression team; the Mouse Facility and the DNA sequencing service at Cogentech. We thank Pier Giuseppe Pelicci and all the participant to the Single-Cell Technoshot (1.0, 2.0) by IEO for support and discussion; Alvis Brazma for insightful discussions throughout the project; Marioni group, Papatheodorou group, Nicassio group for discussions; Marco Cosentino Lagomarsino and Dafne Di Campigli di Giammartino for critical reading the manuscript. Experiments involving animals were carried out in accordance with the Italian Laws (D.lgs. 26/2014), which enforces Dir. 2010/63/EU (Directive 2010/63/EU of the European Parliament and of the Council of 22 September 2010 on the protection of animals used for scientific purposes) and authorized by the Italian Minister of Health with projects 779/2020-PR.

## AUTHOR CONTRIBUTION

FNA: Conceptualization, Methodology, Investigation, Writing – Original Draft Preparation. MJM: Conceptualization, Investigation, Writing – Review and Editing. MPP: Investigation, Writing – Review & Editing. PF: Investigation, Writing – Review & Editing. SP: Formal Analysis. MC: Investigation. PB: Investigation. CR: Resources. BG: Resources. IP: Supervision, Writing – Review & Editing. JCM: Supervision, Writing – Review & Editing. FNI: Conceptualization, Funding Acquisition, Supervision, Writing – Original Draft Preparation.

## DATA AVAILABILITY

The single-cell sequencing data generated in this study have been deposited on ArrayExpress with accession codes E-MTAB-13064 and E-MTAB-13066. The DNA and RNA sequencing data have been deposited on GEO with accession code GSE222596. Whole-exome sequencing data have been deposited on SRA with BioProject ID PRJNA922938.

## CODE AVAILABILITY

The code used to generate the analyses reported in this article will be made freely available at the date of publication.

## METHODS

### Mice

Animals were bred and kept under specific pathogen-free conditions at the Cogentech Mouse Facility located at IFOM (The FIRC Institute of Molecular Oncology, Milan, Italy). All mice were maintained in a controlled environment, at 18-23 °C, 40-60% humidity and with 12-h dark/12-h light cycles, in a certified animal facility. Experiments were performed in compliance with the Italian Minister of Health, the Italian law (D.lgs. 26/2014) and under the control of the institutional organism for animal welfare and ethical approach to animals in experimental procedures (Cogentech OPBA).

### In vivo xenograft

Immunodeficient NOD.Cg-PrkdcscidIL2rgtm1Wjl/SzJ (Also Known As NOD/SCID/IL2Rγ_c_^-/-^) mice were anesthetised by intraperitoneal injections of 1.25% solution of tribromoethanol (0.02 ml/gr of body weight). Barcode-bearing SUM159PT cells were resuspended in a mix of 14 ul PBS amd 6 μl Matrigel and implanted in the fourth inguinal mammary gland of 10-week-old animals. Mice were monitored twice a week by an investigator. For the chemotherapy treatment studies, tumours were allowed to grow to palpable lesions (∼20-30 mm^3^), then mice were randomised into groups and each group treated intraperitoneally with either Paclitaxel (PTX) (10 mg/kg in PBS) or vehicle (PBS) every 5 days for a total of 4 injections. Once the tumours reached 1.2 cm on its largest diameter the animals were euthanized. Tumour growth dynamics was monitored every 3 days by calipers measurements. For in vivo limiting dilution assay (LDA) transplantation experiments, decreasing concentrations (1:10000; 1:1000; 1:100) of SUM159PT were resuspended in a mix of 14 μl PBS and 6 μl Matrigel and transplanted in the fourth inguinal mammary gland of 10-week-old animals. Animals were monitored as before and euthanized after 1.5–3 months (depending on tumour latency).

### Tissue harvest and processing

The primary tumours were removed when the tumour reached an approximate diameter of 1.2 cm. The animals were anesthetised with tribromoethanol, and the tumours were resected. The solid tissue was rinsed with PBS, minced with scalpels, followed by mechanical dissociation using gentleMACS (Mylteni) in digestion mix (collagenase, hyaluronidase, 5 μg ml-1 insulin, Hepes, hydrocortisone). The cell suspension was incubated for 25 min at 37°C followed by an additional step of gentleMACS dissociation. Following a wash with base medium the cells were consecutively passed through 100, 70 and 40 um filters. Primary tumour cells were treated with ACK lysis buffer (Lonza) followed by resuspension in 1% BSA/PBS and processed using the Mouse and Dead Cell depletion kits according to the manufacturer directions (Mylteni).

### Cell cultures

We maintained HEK293T cells in Dulbecco’s Modified Eagle’s Medium (DMEM) with 10% of TET-FREE fetal bovine serum (FBS) and 1% penicillin-streptomycin. Cells were grown in a humidified atmosphere at 5% CO2 at 37°C. SUM159PT cell line and derivatives (Asterand) were cultured in Ham’s F12 medium with 5% TET-FREE fetal bovine serum (FBS), human insulin (5 μg/ml), hydrocortisone (1 μg/ml), and Hepes (10 mM). Barcoded SUM159PT and CRISPRi cell lines medium was supplemented with 2 μg/ml Puromycin for selection. Cells were grown in a humidified atmosphere at 10% CO_2_ at 37°C.

### Perturb_SUM159PT cell line generation

Perturb-seq GBC library (Adamson et al. 2016, Dixit et al. 2016) was a gift from Jonathan Weissman (Addgene ID #85968). The library contains a random 18-nt guide barcode (GBC) close to the polyadenylation signal of the blue fluorescent protein (BFP). The estimated complexity of the library is >5 million unique GBCs. We amplified the Perturb-Seq GBC library in E. coli ElectroMAXTM DH5α-ETM electro-competent cells (Thermo Fisher Scientific), as indicated by the authors (Adamson et al. 2016). DNA extraction was performed with NucleoBond Xtra Maxi (Macherey-Nagel) kit according to the manufacturer’s instructions. Viruses were produced in HEK293T at 80% of confluency with the following transfection mix: 20μg Transfer Vector, 13μg of psPAX2 (gag&pol)(Addgene #12260), 7μg of pMD2G (envelope)(Addgene #12259), 94μL of RT CaCl_2_ and water up to 750μL. Then, 750μL of 2xHBS were added dropwise and 500μL / 10cm plate of transfection mix was added to the cells. The medium was changed after 6h and viruses harvested after 48h, filtered (0.22μm) and ultracentrifuged for 2h at 20000-24000 rpm, at 4°C. The viral pellet was resuspended in 300μL of PBS 1X and stored at –80°C. To produce the Perturb SUM159PT cell line for lineage tracing experiments, 75000 cells were seeded and infected with an estimated multiplicity of infection (MOI)=0.1 in the presence of 8μg/ml of Polybrene (Sigma-Aldrich). After selection, transduction efficiency was measured by FACS analysis, which revealed that 8.6% of cells were successfully infected.

### SUM159PT_KRAB cell line generation and CRISPRi experiments

For the generation of the PB-TRE-dCas9-KRAB plasmid, the DNA sequence of KRAB repressor domain was amplified by PCR from the pHAGE TRE dCas9-KRAB (Addgene plasmid #50917) and cloned in frame into the PB-TRE-dCas9-VPR backbone (Addgene plasmid #63800) within the AscI/AgeI sites. The cloning was sequence-verified by Sanger Sequencing. SUM159PT cells were transfected in MW6 plates, following Lipo3000 transfection protocol (ThermoFisher Scientific) with 500ng of transposon DNA (PB-TRE-dCas9-KRAB) and 200 ng of SuperPiggyBac transposase helper plasmid (SystemsBioscience). After at least 72h from transfection, cells were selected with 200 μg/mL Hygromycin B. The PB-TRE-dCas9-KRAB SUM159PT cell line is referred in the text as SUM159PT_KRAB. We expressed sgRNAs upon cloning into lentiGuide-Puro sgRNA backbone (Addgene #52963) within BsmBI (Esp3I) sites. Lentiviruses were generated in six-well plates, following the Lipofectamine 3000 (Thermo Fisher) protocol. The transfection was performed mixing the construct of interest, psPAX2 (gag&pol)(Addgene #12260) and pCMV-VSV-G (envelope)(Addgene #8454) plasmids at a ratio of 4:3:1. Viruses were collected after 24 h by filtered and frozen. We generated stable cell lines expressing single sgRNAs by lentiviral infection of 150000/well SUM159PT_KRAB cells. Lentiviral supernatants (1:3 dilution) were added to cells, supplemented with 1 μg/mL of polybrene. After 24 hours, cells were selected with 2 μg/ml puromycin (Gibco). For the CRISPRi experiment, we plated sgRNA-expressing stable cell lines with 100ng/mL of doxycycline and harvested cells after 3 days.

### Paclitaxel treatment

A stock aliquot of Paclitaxel (obtained from the IEO hospital) was prepared (70μM) in PBS and used to treat cells. The dose of Paclitaxel used for *in vitro* experiments was established by a dose–response curve of SUM159PT treated for 72 h. IC50 concentrations were estimated by parallel fit estimation (JMP software, *n* = 4). For short-term, single-treatment experiments, Paclitaxel was used at the final concentration of 50nM (∼IC95). After 3 days of treatment, the medium was changed every 3 days without adding the drug. For the generation of the drug resistant model (Figure 5D), SUM159PT cells were treated multiple times with increasing doses of Paclitaxel (10nM, 20nM, 50nM and 100nM). Drug adapted cells were able to survive even when treated with 100nM Paclitaxel and were collected 90 days after treatment for WES analysis.

### TM4SF1^high^ Flow cytometry

SUM159PT_KRAB cells were stained with anti-TM4SF1-APC (clone: REA851 Miltenyi Biotec) antibody for 10’ at 4°C, in the dark and sorted using Fusion Aria Sorter. The bulk population was FAC-sorted using FSC/SSC gate, while the TM4SF1^high^ subpopulation was sorted gating the top 5% APC fluorescence intensity. After passive propagation in vitro, 9 days after sorting, both populations were transplanted into NOD/SCID/IL2Rγ_c_, mice and tumours were collected as described above.

#### RNA sequencing

RNA was extracted from mouse-depleted and dead-cell depleted samples. The bulk and TM4SF1^high^ sorted cells were either immediately processed by RNA-seq or passively propagated in vitro for 43 days (corresponding to passage 19). Cells after 9 and 43 days from sorting were also collected and finally processed by RNA-seq. RNA was harvested using Maxwell RSC miRNA Tissue Kit according to the manufacturer’s protocol. Libraries for RNA-seq were prepared from 1 μg of total RNA using Truseq Total RNA Library Prep Kit (Illumina) following manufacturer’s protocols. Samples were sequenced paired-end 50 bp on an Illumina Novaseq6000 instrument.

#### Whole Exome Sequencing

Sequencing was performed with the Agilent SureSelect All Exon v5 (experiments shown in **Figure 5C,D**) or v7 (experiments shown in **Figure 5A,B**), as by manufacturers’ instructions. Libraries were sequenced with a coverage > 200X on a Novaseq 6000, with a PE 100 reads mode.

### Multiseq – Single-Cell Time-course for assaying drug tolerance

The single-cell time-course experiment for drug tolerance (shown in **Figure 6**) was designed in order to process all time points at the same time. To do so, we performed an en-reverse experiment (i.e. starting from the last time point, D15). The same batch of cells was used for the entire experiment. Cells were passaged every two days and seeded either for passive culture or Paclitaxel treatment (5 x10^6^ cells in 150mm dishes). After 3 days of treatment (50 nM as described above), the medium was changed every 3 days without adding the drug. At day 15, all Paclitaxel time points were collected together, counted and processed with the Multiseq protocol (McGinnis et al. 2019). Briefly, 500000 cells for each sample (D5, D7, D9, D11, D13, D15) were resuspended in 180μL of PBS and then labelled with 20μL of a reaction mix composed of 2 μM of a sample-specific Barcode, 2 μM of LMO-Anchor (a kind gift of the Gartner Lab) and PBS. After 5’ of incubation in ice, we added 20 μL of reaction mix composed of 2 μM of co-anchor in PBS. After 5’ of incubation we stopped the reaction adding PBS/BSA 1%. We centrifuged and washed twice the cells before resuspending each sample in 125 μL PBS/BSA and pooling. After filtering, 25000 cells were loaded on the Chromium Controller. Multi-seq library was isolated from the amplified cDNA and sequenced at 5000 barcode reads/cell depth.

### RT-qPCR

Total RNA was extracted using Maxwell RSC miRNA Tissue Kit according to the manufacturer’s protocol. 1 μg of total RNAs was reverse-transcribed using ImProm-IITM Reverse Transcription System (Promega) and genes were analysed with Quantifast SYBR green master mix (Qiagen). RPLP0 was used as housekeeper gene. The complete list of primers used in this study is reported in **Suppl. Table 16.**

### GBC library preparation from gDNA

The genomic DNA was extracted from at least one million cells (typically 3 million, coverage 300x) using the NucleoSpin Tissue kit (Macherey-Nagel). To enrich for GBCs, six parallel PCR reactions were performed in a final volume of 50 μL using 200ng of genomic DNA, 0.2 μM of dNTPs mix, 0.5 μM of the following primers: F1: GGGTTTAAACGGGCCCTCTA and R4: GCCTGGAAGGCAGAAACGAC and amplified using Phusion^TM^ High-Fidelity DNA Polymerase at the final concentration of 0.02U/µL (Thermo-Scientific) (coverage 50x-500x), in its 5X Phusion HF buffer according to the following PCR protocol: 1) 98° for 30 s, 2) 98° for 7 s, then 60° for 25 s and 72° for 15 s (for 30 cycles), 3) 72° for 5 minutes.

At the end of the PCR, reactions were pooled and purified using QIaquick PCR purification kit (QIAGEN). Then, the eluted DNA samples were run on a 2% agarose gel and the 280bp band purified using the QIAquick Gel extraction kit (QIAGEN). Illumina libraries were generated from 10 ng of DNA, which was blunt-ended, phosphorylated, and tailed with a single 3’ A. An adapter with a single-base ’T’ overhang was added and the ligation products purified and amplified to enrich for fragments that have adapters on both ends.

Libraries with distinct adapter indexes were multiplexed and sequenced (50 bp paired-end mode) on a Novaseq 6000 sequencer.

### Single-cell library and GBC sublibrary preparation (scRNA or snRNA assays)

Single-cell suspensions (500-1000 cells/μL) were mixed with reverse transcription mix using the 10x Genomics Chromium Single-cell 3’ reagent kit protocol V2 (P0_1 and Paclitaxel time-course Replica_2) or V3.1 (P0_2, Paclitaxel time-course Replica_1) and loaded onto 10x Genomics Single-Cell 3’ Chips (www.10xgenomics.com). Libraries were generated as by manufacturers’ instructions and sequenced on Illumina Novaseq 6000 Sequencing System (with a single or dual indexing format according to the manufacturer’s protocol V2 or V3.1). We aimed at a coverage of 50K reads per cell in each sequencing run. Multiome experiments were performed with the Chromium Single-Cell Multiome ATAC + Gene Expression Reagent Kits (V1). Nuclei suspensions (2000 nuclei/μL) were transposed and loaded onto Chromium Next GEM Chip J Single-Cell. Libraries were generated as per manufacturers’ instructions and sequenced on Illumina NOVAseq 6000, aiming at 50K RNA and 50K ATAC reads per cell. To enrich for GBC reads, in a final volume of 50 μL, we amplified by PCR the Perturb-library cassette from 5ng (of the amplified cDNA (as in (Dixit et al. 2016)) using 0.3 μM of dual-indices primers (forward: 5’:AATGATACGGCGACCACCGAGATCTACACCTCCAAGTTCACACTCTTTCCCTACACGACGCTCTTCCGATCT-3’; reverse 5’-CAAGCAGAAGACGGCATACGAGATCGAAGTATACGTGACTGGAGTTCAGACGTGTGCTCTTCCGATCTTAGCAAACTGGGG CACAAGC-3’) and amplified using Q5 2X master mix (NEB #M0541S) according to the following protocol: 1) 98° for 10 s, 2) 98° for 2 s, then 65° for 5 s and 72° for 10 s (25 cycles), 3) 72° for 1 min. The fragment band of the expected length (350-425bp) was purified using EGEL 2% Power Snap Electrophoresis System (Thermoscientific) and checked at bioanalyzer before sequencing.

### sgRNAs list

CRISPRi sgRNAs sequences targeting the two putative enhancers of COL6A1 and COL6A2 were designed using the web tool CRISPick (Broad Institute). The design region was defined by merging the ATAC module regions with the overlapping H3K27Ac signal from Encode (as shown in **Figure 4I**) and exploiting the CRISPRi Range format for unstructured targeting provided by CRISPick. Inputs were: Enh_1 NC_000021.9:+:46048857-46050195 and Enh2 NC_000021.9:+:46052113-46053945. Selected sgRNAs were named according to the position relative to the start of the design window. We selected sgRNAs to cover the entire design window, choosing the sequences with higher on-target activity and filtering out those with potential off-target complementarity. As negative control, we included three scrambled-sequence-sgRNAs for the CRISPRi experiments selected from the previous genome-wide CRISPRi screening library designed by the Weissman lab (Horlbeck et al. 2016) The sgRNAs chosen to target the promoters of COL6A1 and COL6A2 were also chosen from the same study. The list of sgRNAs sequence is reported in **Suppl. Table 16.**

### Genetic barcode analysis

#### Genetic barcode calling

A genetic barcode (GBC) library is built for each sample separately, starting from the FASTQ files, in two steps. First, the 18nt-long sequences located in the GBC locus are extracted using `seqkit amplicon` command from seqkit v2.1.0 (Shen et al. 2016), with either 23-40nt-(DNA reads) or 12-29nt-long (cDNA reads) flanking regions and allowing 1 mismatch. A set *S* of sequences of length 18 is obtained and each *s* ∈ *S* is assigned a weight *w(s)*, corresponding to its frequency in the FASTQ file. Note that, since the relative GBC abundance in a sample is unknown, not accounting for sequencing errors can result in an inflated estimate of sample clonality. Only sequencing errors in the form of mismatches are considered. The underlying assumption is that, if *s,s’* ∈ *S* are sequenced from the same GBC species and carry *d* and *d’* mismatches, respectively, and that *d* < *d’*, then *w(s) ≥ w(s’).* Thus, each GBC species *i* is associated with a subset *S_i_* ⊆ *S*, where the true GBC sequence is the *s* ∈ *S_i_* with maximal weight. The frequency of *i* is the cumulative weight of the sequences in *S_i_*. We infer the set of “true” GBC sequences and simultaneously correct for sequencing errors with the following procedure. Let *G = (V,E,w)* be an undirected graph, where *V = S*, *E* is the set of edges connecting sequences at Hamming distance ≤ *D*, and *w(s)* is the frequency of *s*. We iteratively detect a collection of disjoint subgraphs *G_i_ = (S_i_,E_i_)* induced by *S_i_,* called *stars*, where a node *s* ∈ *S_i_* is the hub and all the other nodes are neighbours of *s*. Stars are defined according to the following conditions: (i) the hub *s* has maximum weight *w(s)* in *S_i_,* (ii) the cumulative weight across stars is maximal, and (iii) the fraction of neighbours of *s* in *G* that do not belong to *S_i_* is < *f*, where 0 ≤ *f <* 1 is a fixed parameter. We compute a heuristic solution with a greedy approach. First, nodes are ordered by non-increasing weight. At each iteration, a new star is created, whose hub is the first node in the list and the other nodes are its neighbours, and the included nodes are removed from the list. The procedure ends when the first star that violates (iii) is found. We set *D* = 1 and *f* = 0.2. The final set of true GBC sequences is defined as the collection of detected hubs and their frequency is the cumulative weight across stars. We approximate the clone content of bulk DNA-Seq sequencing samples as the set of GBCs and their associated frequencies.

#### Definition of tumour initiating clones and drug tolerant clones

In bulk DNA-Seq samples, we approximate clone content with GBC content and clone abundance with count per million reads (CPM). Clones whose frequency significantly differs between conditions are determined as follows. For each clone, we test if the CPM in the condition sample (treated or untreated) is significantly greater compared to the average across control samples (parental), using Fisher’s exact test. The universe is defined as the union of all clones across control and condition samples. P-values are adjusted using the Bonferroni correction method. Clones with adjusted p-value < 0.05 in at least 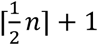 condition samples are labelled *tumour initiating clones* (TICs, if condition is “untreated tumour”) or *drug tolerant clones* (DTCs, if condition is “treated sample” or “treated tumour”), where *n* is the number of condition samples.

### Single-cell RNA-Seq data analysis

#### Cell barcode calling

A GBC reference is defined as the union of the GBC species obtained from cDNA reads, as described in “Genetic barcode calling”, across all samples in the same experiment. Read alignment, UMI counting, cell-containing barcode (CB) calling, and GBC counting are performed using `cellranger count` from CellRanger v6.0 on the human reference genome GRCh38 v2020-A and the GBC reference. Parameter `--expect-cells` is set to 7500 (T0), 20000 (T1 and exp1), and 3000 (exp2). Feature-CB count matrices are obtained, where features denote either genes or GBC species, and entries are UMI counts. Cellular barcodes with less than 5000 (T1 and exp1) or 10000 (T0, exp2) gene UMIs are filtered out. Data post-processing is done with R (R Core Team 2022).

#### MULTI-Seq sample demultiplexing

Samples belonging to T1 and exp1 are demultiplexed using MULTI-Seq barcodes (MBCs), as follows. MBC-containing reads are aligned to the reference barcode sequences using the R package deMULTIplex v1.0.2 (McGinnis et al. 2019). MBC-CB count matrices are obtained. Each MBC univocally identifies a sample, and a sample can be labelled with multiple MBCs. The automatic quantile-based thresholding procedure implemented in deMULTIplex fails to assign a unique MBC label to most of cellular barcodes in exp1, thus a custom demultiplexing procedure is used for both experiments. First, all MBCs with UMI count < 2 are removed from every CB. For every cellular barcode *i*, an MBC *j* is marked as detected if *c_j_* ≥ 15, *c_j_/c_max_* ≥ 0.5, and *p* < 1e-5, where *c_j_* is the UMI count of *j* in *i*, *c_max_* is the UMI count of the top abundant MBC in *i*, and *p* is the probability of observing a value equal or greater than *c_j_* given a Poisson distribution with λ = average UMI count of *j* across cellular barcodes in the sample. Only cellular barcodes with exactly one detected MBC are retained and assigned to the corresponding sample.

#### Clone detection

Expressed GBCs in CBs are identified using the same procedure applied to MBCs and described in “MULTI-Seq sample demultiplexing”, using *c_j_* ≥ 10 and *c_j_/c_max_* ≥ 0.3. A p-value threshold is set only for samples d11-13-15 for exp1 (*p* < 1e-5) and sample d17 for exp2 (*p* < 1e-10). Note that GBC frequency at earlier time points is very low, hence the p-value criterion has no effect. CBs can be assigned 0, 1, or > 1 GBCs, the latter being an effect of multiple cell encapsulation (doublets) or multiple GBCs infecting the same cell (co-infection). We extract clone information from multi-GBC CBs by distinguishing co-infection and doublet events. High UMI doublets are removed using a sample-specific cut-off. Assuming that GBC expression is approximately constant across GBC species and across cells, differences in GBC UMI count are essentially due to droplet-specific mRNA capture. Moreover, the transcriptome size of a cell line model should be constant as well. We deduce that a single cell infected with *k* GBCs should display a higher GBC UMI fraction compared to that of multiple cells encapsulated in the same droplet that jointly account for *k* infection events. For simplicity, GBC species for multiple infection events within the same droplet are assumed to be pairwise distinct. The procedure works as follows. First, each CB set with *k* expressed GBCs is sorted by non-increasing values of *c_i_/u_i_*, where *c_i_* and *u_i_*are the GBC UMI count and the gene UMI count for CB *i*, respectively. We obtain a list of sets *S_2_*,…,*S_m_*, where *m* is the maximum number of distinct GBC species expressed in a CB. Then, we iteratively classify the CBs of each set *S_k_* separately, starting from the smallest *k*. A CB *i* is labelled as co-infection in two cases: (a) all GBCs expressed by *i* are also expressed in at least 5 CBs, including *i*, or (b) the “doublet probability” of *i* (that is, the cumulative sample frequency of each single GBCs expressed in *i*) is below a sample-defined threshold. If neither (a) nor (b) hold, *i* is labelled as doublet. The procedure continues until the fraction of co-infection events in multi-GBC CBs is ≥ 1-*D*, where *D* is the expected doublet fraction in multi-GBC CBs. *D* is a sample-specific doublet rate estimate based on 10x guidelines and the number of called CBs. The clone pool of a single-cell sample is defined as the collection of single GBCs that occur in single infection and doublet events with *k* = 2 GBCs, plus all multi-GBC sets from co-infection events.

#### Clone comparison between single-cell and bulk samples

To assess the accuracy of clonality estimates from bulk samples, we compare the GBC species content between bulk DNA and single-cell RNA GBC sequencing samples from the same condition. We define the frequency of a GBC species in a bulk sample by CPM. Then, for a given pair (*X_b_,X_s_*) of bulk and single-cell samples, we define the value *y = f(x)* as the fraction of GBC species with frequency ≥ *x* in *X_b_* found as expressed in at least one cell of *X_s_*. In practice, we compute *f(x)* in steps of 20 CPM units. Consistent clone estimates in bulk and single-cell samples in a condition result in a monotonically non-decreasing trend of *f(·).* A cell *c* is classified as tumour-initiating or drug-tolerant (see “Detection of tumour initiating clones and drug tolerance clones” above) if and only if all GBC expressed in *c* are tumour-initiating or drug-tolerant, respectively. Conversely, the single-cell cluster labelling is transferred to a bulk sample as follows: the GBC abundance (CPM) in the bulk sample is first normalised by its average abundance at baseline; then, the abundance of cluster *C* in the bulk sample is calculated as the contribution of all GBCs expressed by any cell in *C*.

#### Quality filtering and normalisation

All CBs with no expressed GBCs or classified as doublets are removed from subsequent analysis. Samples from technical replicates (same vial) are concatenated and processed using Seurat v4.0.5 (Hao et al. 2021). UMI counts are added a pseudocount = 1, divided by library size, multiplied by 10000, and log-transformed (natural logarithm), to obtain log-normalised counts.

#### Dimensional reduction and clustering

Highly variable genes detection is performed on log-normalised counts using two different approaches: `min.var.plot` and `vst.` We set `xmin` = 0.1 and `xmax` = 10 for `min.var.plot`. The input parameters for each algorithm are let vary in a pre-defined set: dispersion.cutoff in {0.5,1,1,5} for `min.var.plot`, and `nfeatures` in {1000,2000,5000} for `vst`. A cell-cycle score is computed using the Seurat function `CellCycleScoring` with default parameters. We observe a high cell-cycle effect in parental samples (T0 and T1); hence, the cell-cycle score computed as above is regressed out before scaling and centering. PCA is performed on z-scores of the log-normalised counts on the reduced space of highly variable genes. A total of 50 PCs is computed. The optimal number *n* of PCs to retain for clustering is defined as the minimum *n* such that the standard deviation explained by the *n*-th PC exceeds 50% of the average across the 40th to 50th PCs. Multiple Louvain clustering runs are performed on the selected PCs, for each highly variable gene set, by varying the number of neighbours *k* in the knn graph and the resolution parameter *r* in a pre-defined set: *k* ∈ {30,40,50} and *r* from 0.1 to 0.8 in steps of 0.1. A second and third round of highly variable gene selection, PCA, and clustering are possibly performed after the removal of small clusters with very low UMI count, found in specific solutions, until the average UMI count is homogeneous across clusters. We redo the whole clustering procedure instead of just removing the small, low-quality clusters, because they are usually outliers in the expression profile of the sample and can affect the definition of highly variable genes. We obtain 9395 and 13562 cells for T0 and T1, and 7884 and 10698 cells for exp1 and exp2, respectively. For T0 and T1, a silhouette score is computed for each clustering solution on the euclidean distances calculated on the same PCs used for clustering. The clustering solution with the highest silhouette score is defined as optimal. To increase the number of detected clusters, we set a higher value for the resolution parameter, while maintaining the other parameters of the optimal solution unchanged (the highly variable gene set and the number of neighbours in the knn graph). The final clustering solutions for T0 and T1 are obtained with `vst` method by setting `nfeatures` = 1000 and clustering parameters *k* = 40 and *r* = 0.5, resulting in 7 clusters for both experiments. For exp1, we obtain 3 clusters using the following settings: `vst` method, `nfeatures` = 5000, *k* = 30, and *r* = 0.1. UMAP reduction (Leland McInnes 2018) is run with RunUMAP using default parameters on the same input used for clustering.

#### Differential expression analysis and functional annotation

A differential expression analysis is run between conditions via `FindMarkers` function from Seurat with MAST method (Yajima 2022)and accounting for sample ID as a covariate. We only test for genes detected (UMI count > 0) in at least 10% cells in either condition such that |log_2_(FC)| ≥ 0.25. A gene is defined as differentially expressed if its adjusted p-value is < 0.05 (Bonferroni correction) between the two conditions.

#### Computation of cell-cell distance distributions

To evaluate the relationship between clonality and gene expression at basal state, we compare sister cells (same clone) and non-sister cells (different clones) at T0 and T1. We note that a sister cell similarity measure within the same experiment would be biassed towards clones with vial-specific frequency > 1, thus we opt for a comparison between experiments (1886 common clones). Cells of T0 and T1 are projected on a common latent space using with Canonical Correlation Analysis (CCA, (Butler et al. 2018), using the intersection of highly variable gene sets (defined as in “Dimensional reduction and clustering”) between experiments. By definition, this integration approach removes dataset-specific complexity by keeping only shared correlations, hence the contribution of the experiment covariate to cell-cell similarity should be minimised compared to more conservative approaches. Inter-sample cell-cell Euclidean distances are computed on the CCA latent space on sister cell pairs and a random subset of non-sister pairs of the same size. A gaussian kernel density estimation is computed for sister and non-sister distances separately to obtain a global distance distribution. Then, to evaluate the relationship between and within the DTC and the non-DTC pool before and after treatment, we computed the Euclidean distances between cells at T1 (baseline) and in exp1 (after treatment); finally, we evaluated the relationship between clonality and gene expression on the treated condition by computing intra-sample Euclidean distances at day ≥ 13 on the PC space of exp1 and exp2, as defined in “Dimensional reduction and clustering”. We obtained Gaussian kernel density estimates for both cell-cell distance matrices as above.

#### Computation of clone sharedness and detection of cell subpopulations

To assess whether sister cell similarity is uniform across clones or is rather clone-specific, we introduce a *clone sharedness score* across clusters (see “Dimensional reduction and clustering”). It is defined, for each pair of clusters *i* and *j* in T0 and T1, as *cs*(*i,j*) = (*n_ij_·n’_ji_*)/(*N_i_·N’_j_*), where *n_ij_* (*n’_ji_*) is the number of cells in *i* (in *j*) with at least one sister in *j* (in *i*) and *N_i_* (*N’_j_*) is the total number of cells in *i* (in *j*). Each cluster *i* in T0 is matched with the cluster *j*^max^ in T1 with maximum clone sharedness with i, namely, *j*^max^ = arg max*_j_ cs*(*i,j*). The three clusters belonging to maximal clone sharedness pairs (*i,j*^max^) are defined as subpopulations and labelled by non-increasing value of *cs*(*i,j*^max^) as S1, S2, and S3. Transcriptionally stable clones are defined as the ones only found in one of the three subpopulations in both T0 and T1, they are 437 in total. This approach is independent of single-cell data integration, whose accuracy is difficult to assess when reliable markers are unknown (Tran et al. 2020, Luecken et al. 2022). We define the gene signature of *S* ∈ {S1,S2,S3} as the set of genes that are differentially upregulated between *S* and its complement in both T0 and T1. We obtain 109, 29, and 27 genes for signatures S1, S2, and S3.

#### Definition of clone-clone pair propensity

We define a *pair propensity* score to measure the pairwise gene expression similarity bias between clones. For each cell, we consider its directed neighbourhood of size *k*, *i.e.*, each cell has exactly *k* nearest neighbours and the neighbour relationship is not symmetric. Given two clones labels *i* and *j*, possibly identical, the observed (*i,j*) frequency *f ^obs^* is the number of cell pairs (*c,c*) such that *c* and *c* belong to clones *i* and *j*, respectively, and *c_j_* is a neighbour of *c_i_*. The expected (*i,j*) frequency *f ^exp^*is calculated based on the frequency of clones *i* and *j* and the neighbourhood size *k*, as follows. Given the neighbourhood of *c_i_,* the probability that a given cell *c* ≠ *c_i_* in the neighbourhood is labelled with *j* is given by *(n_j_-I(i,j))/(N-1)*, where *n_j_* is the number of cells labelled with *j*, *N* is the total number of cells in the sample, and *I(i,j)* is the indicator function (*I(i,j)* = 1 if *i* = *j* and *I(i,j)* = 0 if *i* ≠ *j*). We obtain *f ^exp^* = *n ·k·(n -I(i,j))/(N-1)*, where *n*_i_ is the number of cells labelled with clone *i*. The pair propensity of *i* and *j* is defined as *p_ij_* = *f ^obs^*/*f ^exp^*. By iterating across all clone pairs, we obtain a non-symmetric matrix *P* with *n* rows and *n* columns, where *n* is the number of distinct clone labels. The value of *p_ij_* indicates the propensity of clones *i* and *j* to be closer (*p_ij_* > 1) or farther (*p_ij_* < 1) from one another in the sample than expected by chance, where *p_ij_* = 1 denotes random association. Finally, we obtain a symmetric matrix 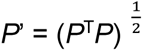.

#### Detection of clone lineages

We use the above pair propensity definition to find groups of clones with mutually similar gene expression profiles on the treated condition at day ≥ 13. We first define the neighbour relationship, as follows. The top 1000 highly variable genes are computed with `vst` method on the union of the time points in exp1 and exp2, respectively. For each sample, a knn graph with *k* = 40 is built on the top *n* PCs (see “Dimensional reduction and clustering”). Matrix *P’* (see “Definition of clone-clone pair propensity”) is computed on the top 20 highest frequency clones, clones *i* such that *p_ii_* < 1 are discarded, and the matrix obtained is clustered using the `hclust` function with `ward.D2` method on Euclidean distances. We obtain two clusters *A* and *B* that serve as “anchor” to add the remaining clones. Clones not yet included in *A* nor in *B* are sorted by non-increasing frequency and we iteratively add one clone at a time, as follows. The first clone *i* in the ordering is considered and a pair propensity *p_iA_* (*p_iB_*) is computed between *i* and the union of clones in *A* (in *B*). If *p_iB_* < 1 and *p_iA_* > *p_iB_* + 1, then *i* is added to *A* (likewise for *B*), otherwise it is discarded. The procedure continues until the last clone in the ordering is considered. Starting from the sets *A* and *B* computed on each of the four samples, we define lineage L1 (lineage L2) as the set of clones always assigned to *A* (to *B*) and detected at least once in both exp1 and exp2. We detect 11 clones for L1 and 17 clones for L2. We define the gene signature of L1 (of L2) as the set of genes that are differentially upregulated (downregulated) between cells in L1 and L2 in both exp1 and exp2. We obtain 42 and 46 genes for L1 and L2.

#### Functional annotation

We perform pathway enrichment analysis (REACTOME (Gillespie et al. 2022)) as follows. Genes with official gene symbols are first converted to ENTREZ identifiers, using limma v3.49.5 (Ritchie et al. 2015) and clusterProfiler v4.1.4 (Wu et al. 2021) packages, respectively. Enrichment significance is calculated with a Fisher’s exact test using the `enrichPathway` function from clusterProfiler. Benjamini-Hochberg method is used to correct for multiple testing. We perform Gene Set Enrichment Analysis (GSEA (Sergushichev 2016)) as follows. Given a query case-control, the list of genes (universe) is ordered by non-increasing log_2_(FC) values; GSEA is run with `GSEA` function from clusterProfiler, with `nPerm` = 1000 sample permutations. MYC activity was computed with `ModuleScore` function from Seurat on T0 and exp1 cells using HALLMARK_MYC_TARGETS_V1 and HALLMARK_MYC_TARGETS_V2 (www.gsea-msigdb.org) as MYC target gene lists.

### Single-cell multiome data analysis

#### Cell barcode calling

A GBC reference is defined as the union of the GBC sets computed from the scRNA-Seq reads, as described in “Genetic barcode calling”. Read alignment, UMI count, peak calling, and CB calling are performed using `cellranger-arc count` from CellRanger v6.0 on genome reference GRCh38 v2020-A-2.0.0. GBC counts are extracted using `cellranger count`, setting `--expect-cells` to 6200 (M0_1) and 8800 (M0_2) and `--include-introns`. CBs with less than 10000 UMIs are filtered out.

#### Clone detection and quality filtering

Expressed GBCs in CBs are identified using the same procedure applied to scRNA-Seq samples and described in “Clone detection”, using *c_j_* ≥ 10 and *c_j_/c_max_*≥ 0.3, and no p-value threshold. All CBs with no expressed GBCs or classified as doublets are removed from subsequent analysis. We obtain 2446 and 2377 nuclei for M0_1 and M0_2, respectively.

#### Gene expression analysis

Cell clusters are defined using gene expression information only, on M0_1 and M0_2 separately. UMI count normalisation is calculated as for scRNA-Seq datasets with Seurat v4.0.6. Highly variable genes are selected using `FindVariableFeatures` function from with method = `vst` and `nfeatures` = 5000. PCA is performed on the set of highly variable genes using `RunPCA` function with default parameters. The first 30 PCs are used to compute a Shared Nearest Neighbours graph via the `FindNeighbors` function, then `FindClusters` function is used to cluster the cells, using the SLM algorithm with resolution ranging between 0.2 and 1.2 in steps of 0.1. A silhouette width is calculated for each clustering solution on the Euclidean distance matrix computed on the same input used for clustering. Subpopulations S1, S2, and S3 in the two replicates are detected by matching the clusters to T0 and T1 using the sharedness score (see “Computation of clone sharedness and detection of cell subpopulations). The selected resolution value is the one that maximises the average silhouette width while keeping the three subpopulations distinct. We select resolution 0.4 and 0.6 for M0_1 and M0_2, respectively, and obtain 7 and 6 clusters. UMAP projection is calculated using `RunUMAP` with default parameters on the same input used for clustering. Cluster markers are extracted with the Wilcoxon Rank-sum test implemented in `FindAllMarkers` function using default parameters. Differentially expressed genes and gene signatures for each subpopulation are obtained as in “Single-cell RNA-Seq data analysis”. Subpopulation gene signatures are defined as for the scRNA-Seq samples.

#### Chromatin accessibility analysis - Data preprocessing

The region-cell count matrices, where regions are ATAC peaks and entries are fragment counts, are processed as follows. Raw fragment counts are normalised using TF-IDF (Term Frequency - Inverse Document Frequency), which assigns higher importance to highly cell-specific regions. The active regions (fragment count ≥ 1) in at least 10 cells are selected using the `FindTopFeatures` function from Signac v1.7.0 (Stuart et al. 2021). LSI (Latent Semantic Indexing) is applied to reduce the dimensionality of the dataset. We compute 50 LSI dimensions. We keep dimensions from 2 to 50, as dimension 1 shows positive correlation with sequencing depth. UMAP projection is calculated on the same LSI space, using default parameters.

We define pairs of regions associated with the same transposition event between replicates as those lying at a distance ≤ *d* on the genome. `findOverlap` function from GenomicRanges v1.46.1 (Lawrence et al. 2013) is used to determine such pairs, by varying the gap size (*i.e.*, the value of *d*) from -1000 (overlap) to 1000 (padding) in steps of 10. Depending on *d*, each region in M0_1 can have 0, 1 or multiple matching regions with M0_2, and *vice versa*. The idea is that a low *d* would fail to recognise regions stemming from the same transposition event, whereas a high *d* would result in many spurious matches. Low (high) values for *d* result in few (many) multiple matches. The value of *d* is chosen such that both *u_d_* and *u_d_*/(*N_d_*-*u_d_*) are maximised, where *u_d_* is the number of unique matches between M0_1 and M0_2 and *N_d_* the total number of matches. We select *d* = 0 and obtain 120,414 and 120,377 matched regions in M0_1 and M0_2, respectively.

#### Chromatin accessibility analysis - Topic modelling

We use a topic modelling approach to group regions that are consistently open in the same sets of cells while reducing data sparsity. Given a region-cell matrix, topic modelling outputs a set of *topics*, where each region has a probability of being assigned to a topic and each topic has a probability of being assigned to a cell. Modelling chromatin accessibility in this way has three advantages: first, the number of topics is typically orders of magnitude smaller compared to the number of regions; second, a cell’s epigenome is expressed as the contribution of multiple topics; third, the importance of a region in a cell is interpretable, namely, as the combination of the weight of the region in a topic and the contribution of that topic in the cell’s epigenome.

Specifically, we use the Latent Dirichlet allocation (LDA) model implemented in cisTopic v0.3.0 (Bravo Gonzalez-Blas et al. 2019) on each replicate separately, on the set of matching regions between the two replicates (both unique and multiple matches, see “Chromatin accessibility analysis - Data preprocessing”). The input to cisTopic is a raw region-cell count matrix, binarised by setting a cutoff at 1 fragment per region per cell. cisTopic is run using a total number of 1000 iterations and a burn-in period of 500 iterations to the Collapsed Gibbs Sampler. The procedure is repeated by varying the number of output topics *n* between 10 and 50 in steps of 10. We select the maximum log-likelihood solution and obtain *n* = 40 for both M0_1 and M0_2. Topics with Pearson correlation coefficient > 0.5 between topic-cell probability and fragment count are discarded. We select 31 topics for M0_1 and 34 topics for M0_2.

#### Chromatin accessibility analysis - Detection of ATAC modules

To select the topics that represent robust biological signals, we compute a topic reproducibility score between replicates, as follows. To this aim, an Irreproducible Discovery Rate (IDR) (Li et al. 2011) is calculated between every pair of topics in M0_1 and M0_2, respectively, on region-topic probabilities, using idr v2.0.3 tool with parameters `--peak-merge-method max --idr-threshold 0.05 --max-iterations 100`. Given topics *t_1_* and *t_2_* in M0_1 and M0_2 and a region *r*, the IDR statistic expresses the probability that the region-topic probabilities of *r* in *t_1_* and *t_2_*are different. We say that *r* is reproducible between *t_1_*and *t_2_* if IDR < 0.05. For each topic pair (*t_1_,t_2_*) in M0_1 and M0_2, we define the reproducibility score as the weighted mean between the number of reproducible regions and the 75th percentile of min {−125 log_2_(IDR), 1000} across those regions. We define the sets of regions with IDR < 0.05 in topic pairs with reproducibility score > 0 as *ATAC modules*. For each cell and for each module, we compute a *module AUC* (Area Under the Receiver Operating Characteristic curve) score, where the TF-IDF value is the predictor variable and the membership to the module is the response variable, using the `auc` function from pROC v1.18.0 (Robin et al. 2011) and direction “<”. The set of regions resulting from the union of all ATAC modules were annotated using the ENCODE registry of Regulatory Elements v2 (cCRE) (Consortium et al. 2020) with a minimum overlap of 1bp.

#### Relationship between epigenetic modules and subpopulations

We detect the ATAC modules explaining specific cell subpopulations as follows: for each ATAC module, we compute an AUC score where the predictor variable is the module AUC defined above and the response variable is either the subpopulation membership (S1,S2,S3) or the phenotype (TIC,DTC). Then, we compute Spearman’s rho between the module AUC for each of the top 40 reproducible ATAC modules and each gene among those detected in at least 20 cells in the highly variable gene pool (`vst` method, `nfeatures` = 5000) across all nuclei. We annotate each gene according to whether they belong to S1, S2, and S3 signatures (obtained from both scRNA-Seq and multiome gene expression data) or are annotated as human TF by Uniprot(Consortium 2018) (GO:0003700, taxon = human, gene product type = protein; 1435 TF in total). To identify putative cis-regulatory regions, we extract the genes in the transcriptional signatures of S1, S2, and S3 that are proximal to reproducible regions in epigenetic modules, with a neighbourhood size of ±50 kbp around the region, using `findOverlaps` function from GenomicRanges package.

#### Identification of enriched TFs

For a given module, we extract two sets of genes according to the following criteria: (i) Spearman’s rho ≥ 0.2 between the module AUC and gene expression across nuclei (see above), or (ii) gene locus lying ≤ 100 kb away from any region in the module, as by GREAT 4.0.4 (McLean et al. 2010) analysis, using the basal plus extension gene regulatory domain definition. These two sets are separately used as input to ChEA v.3 (Keenan et al. 2019) and the output ranked list of enriched TFs is obtained.

### Whole-exome sequencing data analysis

#### Read alignment, variant calling, and extraction of reproducible CNVs

Reads were aligned with BWA (-t 16) v0.7.17 (Li and Durbin 2009) and copy number variants were called with CNVkit v0.9.8 (Talevich et al. 2016) with default parameters, using the parental SUM159PT cells as a normal reference sample and providing the appropriate Agilent bed file (v5 or v7) as `target`. The chromosomal copy number variants detected with CNVkit were retrieved. CNVs were intersected across replicates using `bedtools intersect` from bedtools v2.30.0 (Quinlan and Hall 2010), requiring ≥ 1 bp overlap. Only regions covered by all replicates are retained. A CNV consensus is then defined as the set of regions that span > 80% of a CNV in each replicate.

#### Comparison with single-cell sequencing data

Regions from scATAC-Seq assays of M0_1 and M0_2 are intersected with CNV consensus regions using subsetByOverlap function from GenomicRanges v1.45.0 with default parameters. Genes from scRNA-Seq assays are intersected with CNV consensus regions using the same approach but allowing ±100 kbp around the CNV consensus regions. Gene loci coordinates are extracted from the annotation used in the single-cell multiome analysis.

### Bulk RNA-Seq analysis of SUM159PT and tumours

Reads were trimmed, filtered and aligned using STAR v2.7.3 (Dobin et al. 2013). Read count extraction and TPM normalization were performed using `FeatureCounts`. The TM4SF1^high^ signature, consisting of 433 differentially expressed genes (DEGs), was defined using edgeR (within Galaxy v3.36), considering the sorting batch as a factor, and selecting genes with log_2_(FC) > 1 and p-value < 0.05 (N=3 independent sorting batches). The functional analyses of TM4SF1^high^ DEGs were generated through the use of IPA (QIAGEN Inc.). Differential gene expression analysis in tumours versus 2D samples was performed with the Bioconductor Deseq2 package v1.34 (Love et al. 2014) using default parameters.

### Comparison with single-cell sequencing datasets from the literature

#### Triple-negative breast cancer (TNBC) scRNA-Seq

Concatenated scRNA-seq data from primary TNBC samples were retrieved from GSE161529 (Pal et al. 2021). Filtered cell-count matrices were processed as in “Quality filtering and normalisation”. Epithelial cells were defined by normalised EPCAM expression greater than the local minimum between the first and second maximum in the distribution. Cells with UMI count > 20000 were removed, resulting in 9063 epithelial cells for downstream analysis. Highly variable gene detection, PCA, and clustering are performed with `Seurat_preprocessing` function from Scissor 2.0.0(Sun et al. 2022). Pseudo-bulk positive and negative phenotypes are generated for each subpopulation (S1,S2,S3) and its complement, respectively, on each scRNA-Seq sample at baseline separately (T0_1, T0_2, T1_1, T1_2), resulting in n = 8 samples per phenotype pair (4 replicates per condition). Scissor is run on the single-cell dataset for each of the 3 phenotype pairs. Parameter `alpha` for the logit regression is let vary in {0.7,0.8,0.9}, until ≤ 15% phenotype-associated cells (both positive and negative) are detected (`cutoff` = 0.15). UMAP is run with default parameters on the top 10 PCs.

#### Pancreatic ductal adenocarcinoma (PDAC) scRNA-Seq

Association with subclonal dissemination for 2010 genes were retrieved from (Simeonov et al. 2021). To map S1 and S3 signature genes, murine gene symbols were converted to human gene symbols with limma v3.49.5.

## Supplementary figure legends

**Figure S1.**
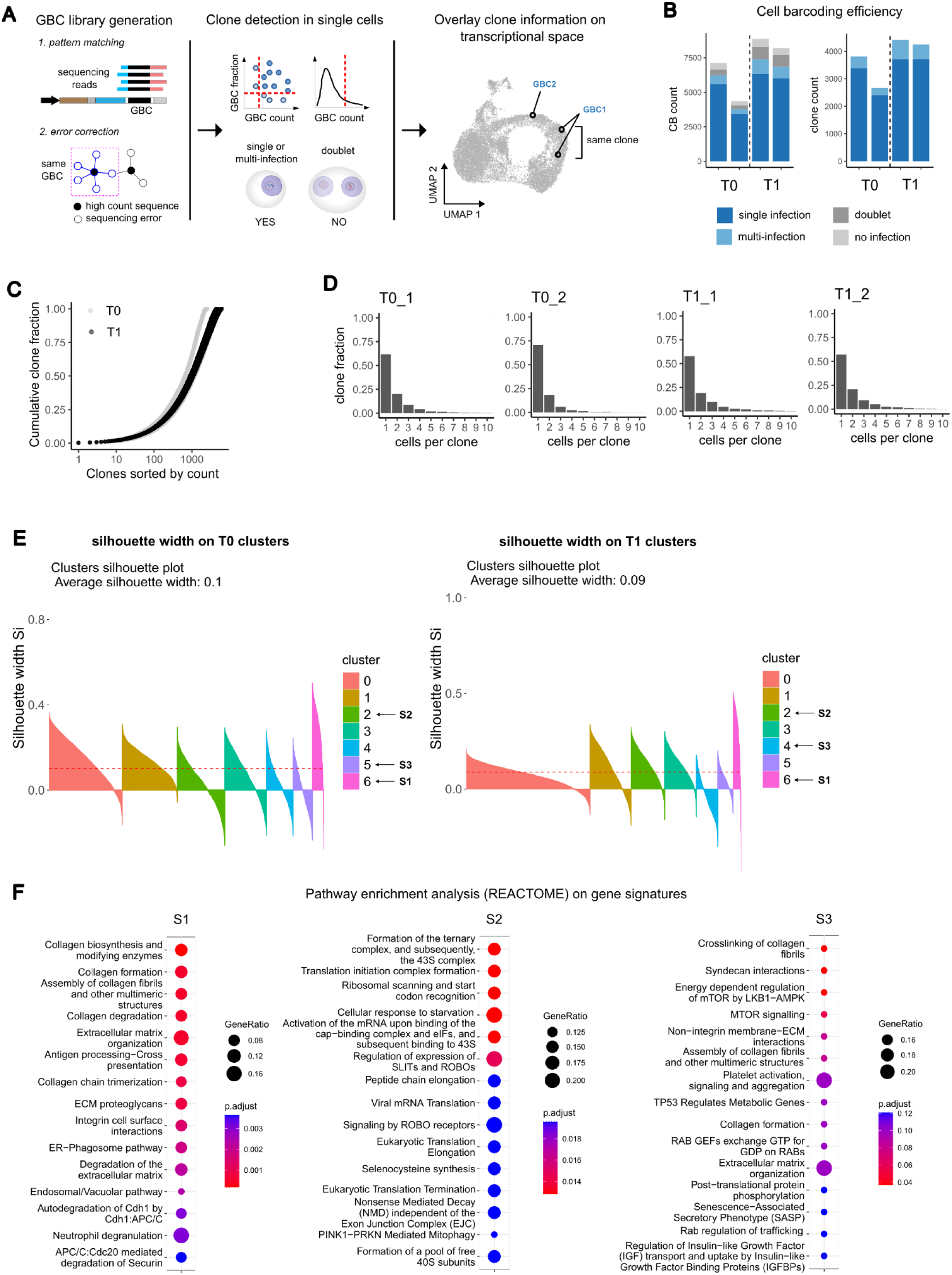
**A.** Left: the GBC sequence library is generated in two analysis steps: 1. sequences on the GBC locus are extracted from the reads via pattern matching; 2. error-correction is performed using a graph approach. Centre: expressed GBCs in single cells are detected, single-cell-containing droplets are selected, and cells are assigned to clones accordingly. Right: sample UMAP representation showing the overlay of clone information on cells in gene expression space. **B.** Clone calling for T0 and T1 (n=2 replicates per condition). Left: cellular barcode (CB) classification into single infection, multi-infection, doublet, and no infection; CB count (left) and clone count (right) are shown. Clones from both single-infection, co-infections, and doublets with exactly 2 expressed GBCs are retained (see Methods). **C.** Comparison of cumulative clone distributions in parental (T0) and untreated (T1) samples. **D.** Relative clone frequency as a function of the number of cells per clone for T0 and T1 in each duplicate. **E.** Silhouette width for each cell in the clustering solutions shown on Figure 1D, for T0 (left) and T1 (right) scRNA-Seq samples (n=2 per time point). **F.** Pathway enrichment analysis (REACTOME) of subpopulation gene signatures. The top 15 significantly enriched terms (q-value < 0.1) are reported and sorted by non-increasing q-value. The size of the circles is proportional to the fraction of genes found in each pathway for each signature.

**Figure S2.**
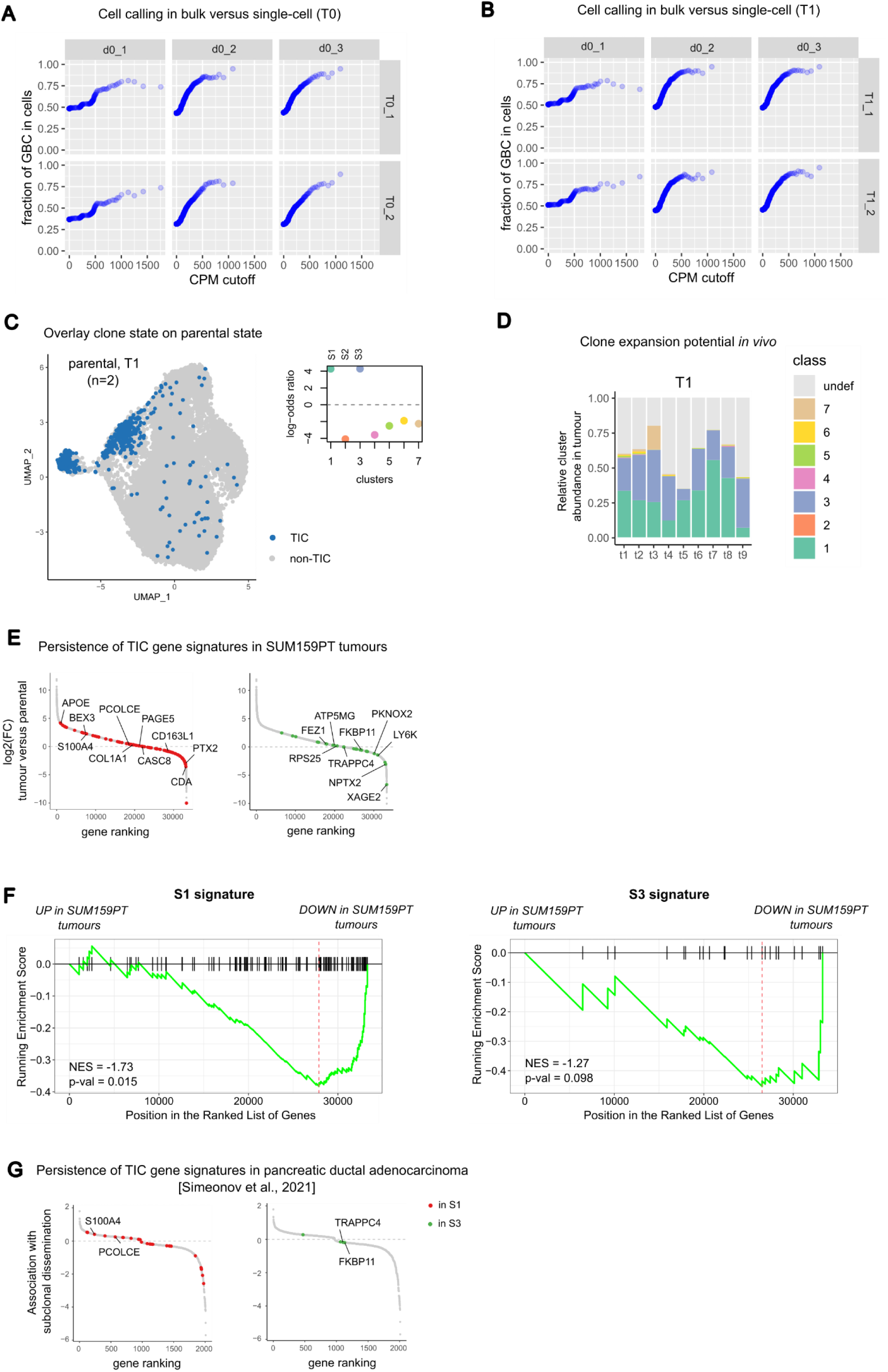
**A.** Comparison of clone calling by single-cell (T0, rows) and clone estimate by bulk DNA sequencing (d0, columns). GBCs are binned according to CPM values (see Methods). The *x* axis represents the midpoint of the intervals corresponding to the bins; the *y* axis represents the fraction of GBCs detected in each scRNA-Seq sample whose abundance in the bulk lies in the interval corresponding to the *x* axis (see Methods). **B.** As in A, with the comparison of clone calling by single-cell (T1, rows) and clone estimate by bulk DNA sequencing (d0, columns). **C.** Mapping of the tumour initiating clones at parental state (T1). Left: UMAP representation of T1 cells on gene expression space; the 477 cells classified as TIC are coloured in blue. Right: the *x* axis is the cluster identifier, and the *y* axis is the log-odds-ratio obtained from the contingency table comparing cluster assignment and TIC labelling across cells at T1. **D.** Association between parental state (T1) and clone expansion *in vivo*. The barplot shows the relative abundance of T1 clusters in every tumour (unassigned clones shown in grey). **E.** Differential expression analysis comparing SUM159PT tumours and the parental population. The *x* axis represents the genes ranked by non-decreasing log_2_(FC) between the average expression in tumours (n=5) and parental samples (n=6); the *y* axis is the log_2_(FC) value; genes in the S1 (S3) signature are marked in red (green) and the top log_2_(FC) genes in the scRNA-Seq assay are labelled. **F.** GSEA of S1 (left) and S3 (right) gene signatures on the list of genes ranked by log_2_(FC) expression in SUM159PT tumours compared to baseline. **G.** S1 and S3 gene signatures in the scRNA-Seq dataset of pancreatic ductal adenocarcinoma from (Pal et al. 2021). The *x* axis represents the genes ranked by non-decreasing association with subclonal dissemination in pancreatic metastatic clones; the *y* axis is the association with subclonal dissemination. Genes in S1 and S3 signatures are coloured as in **E**.

**Figure S3.**
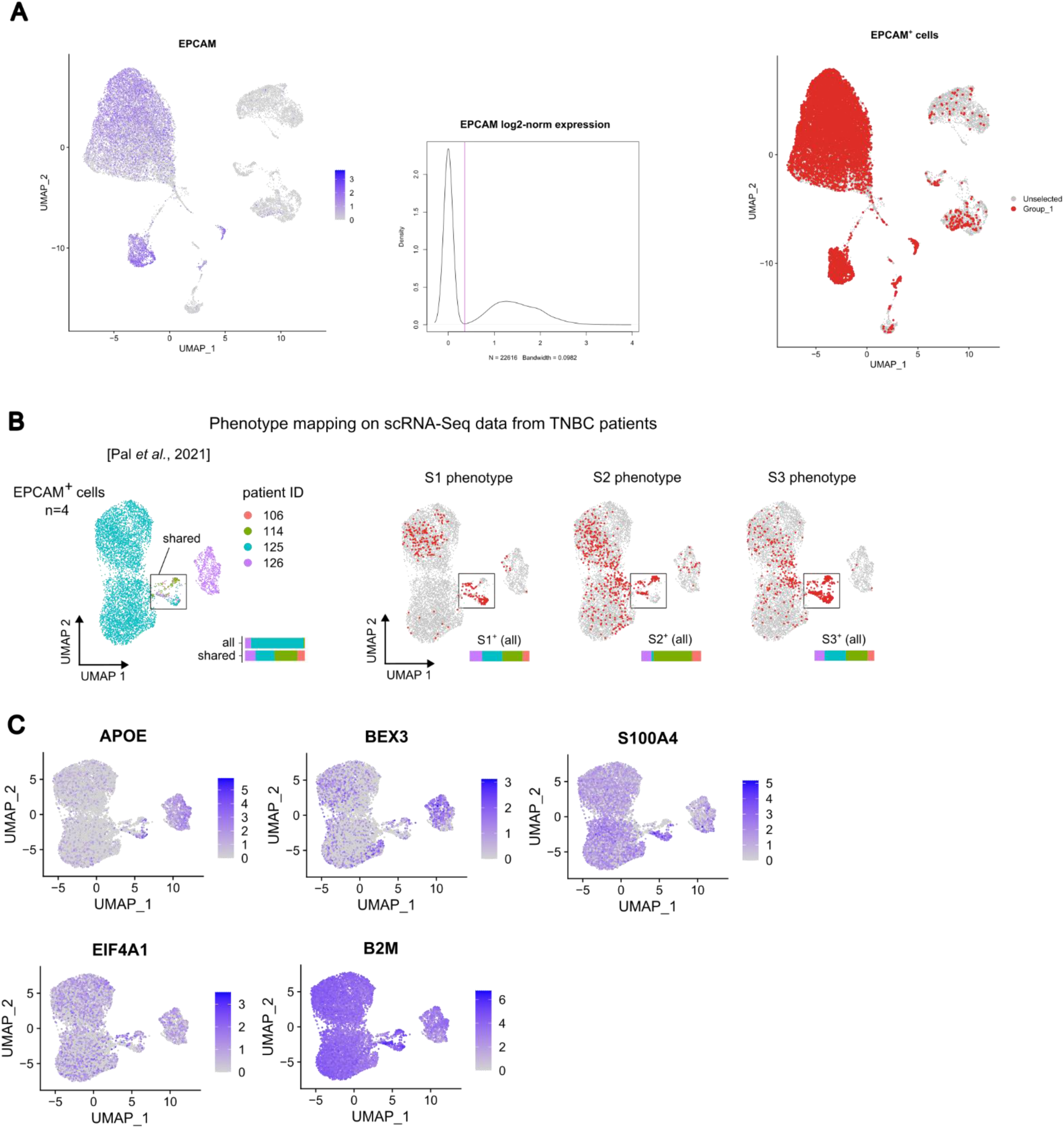
**A.** Selection of epithelial cells from (Pal et al. 2021) dataset. Left: cells on UMAP gene expression space from n=4 primary TNBC samples, coloured according to log-normalised EPCAM expression. Centre: gaussian kernel density of log-normalised EPCAM expression; a vertical line coloured in magenta separates non-epithelial (EPCAM^-^, left mode) and epithelial cells (EPCAM^+^, right mode). Right: cells on UMAP gene expression space where epithelial cells are coloured in red and non-epithelial cells are coloured in grey. **B.** Transcriptional phenotype inference for S1, S2, and S3 subpopulations on (Pal et al. 2021) dataset. Left: the epithelial (EPCAM^+^) cells from n=4 primary TNBC samples, defined as in A., are plotted on gene expression space (UMAP) and coloured by sample (9063 cells in total); the composition of the whole set of cells and of the cluster of cells shared among samples (indicated with a rectangle) is reported with a coloured bar. Right: Scissor output for each phenotype, as defined by bulk S1, S2, and S3 gene expression, on the EPCAM^+^ cells defined above; cells predicted as phenotype-positive are highlighted in red; the sample composition of phenotype-positive cells is reported with a coloured bar. **C.** Epithelial cells plotted on gene expression space (UMAP), as in **B**, coloured according to the log-normalised expression of S1 signature genes; the reported genes are significantly upregulated both in S1^+^ cells (compared to S1 complement) and in SUM159PT tumours (compared to baseline SUM159PT expression).

**Figure S4.**
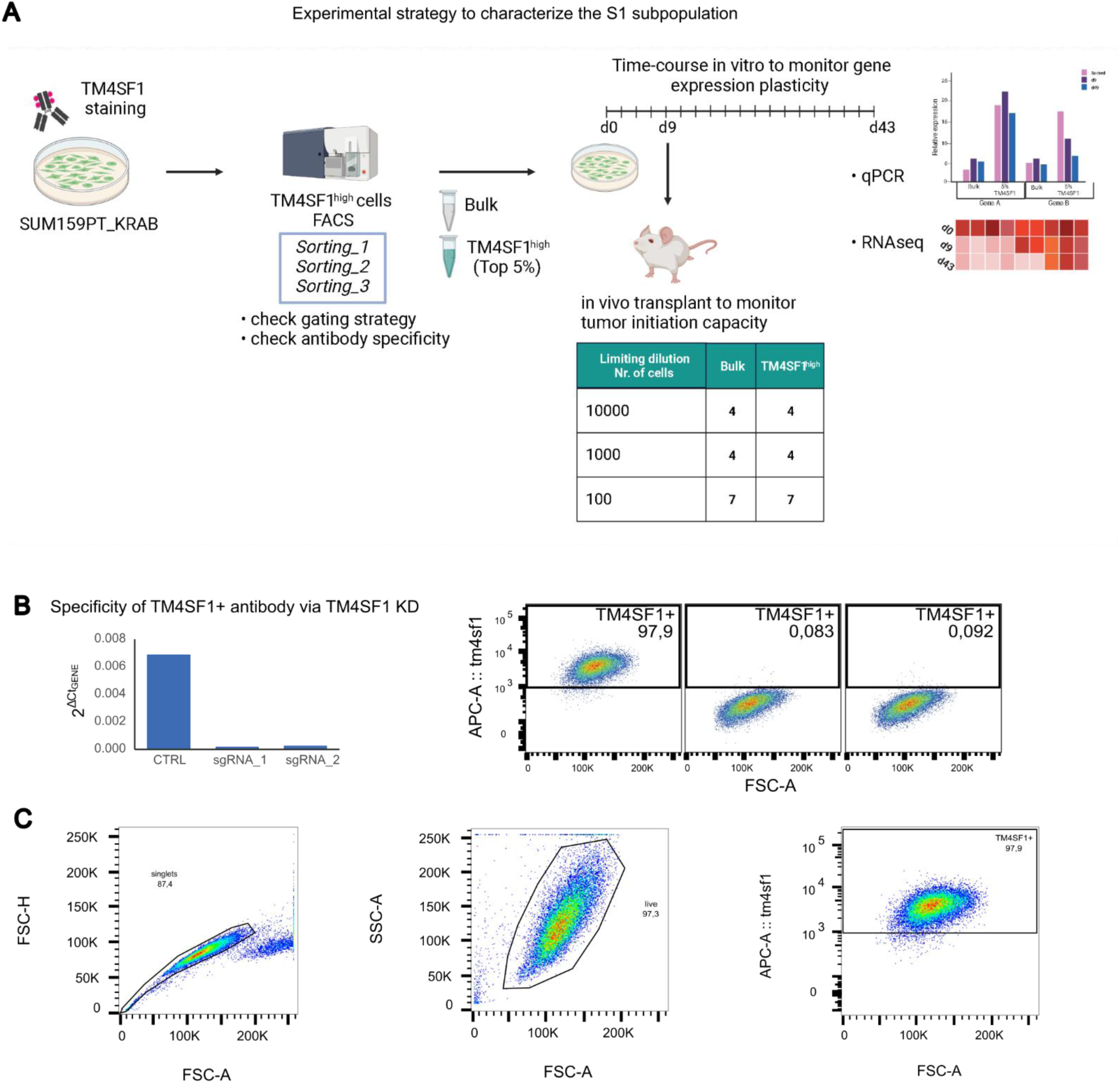
**A.** Experimental design to characterise the S1 subpopulation: SUM159PT (expressing a doxycycline inducible dCas9-KRAB transgene, SUM159PT_KRAB hereafter) were stained with an APC-conjugated TM4SF1 antibody. Bulk and TM4SF1^high^ subpopulations were FAC-sorted (n=3) and processed for RNA extraction and subsequent RT-qPCR and RNA-seq analysis. For the time-course in vitro experiment, the two subpopulations were grown for 43 days and pellets were collected at every passage. RT-qPCR was performed on time points every 3-4 passages (n=7). The limiting dilution transplantation experiment *in vivo* was performed using frozen vials of bulk and TM4SF1^high^ cells passage 2 (P2) and propagated in 2D for other two passages (day 9 from sorting). Cells were first checked for subpopulation gene signature expression by qPCR, see **Figure S5A**). **B.** Left: RT-qPCR data reporting the expression of TM4SF1 in SUM159PT_KRAB cells infected with two sgRNAs targeting TM4SF1 or a control sgRNA (CTRL) and induced for 72h with doxycycline. Expression levels are normalized to the expression of the housekeeper gene RPLP0. Right: FACS plots representing the percentage of TM4SF1+ cells in control and sgTM4SF1 after 72h of doxycycline treatment and staining with the APC-conjugated TM4SF1 antibody. **C.** Gating strategy for the control experiment in **B**.

**Figure S5.**
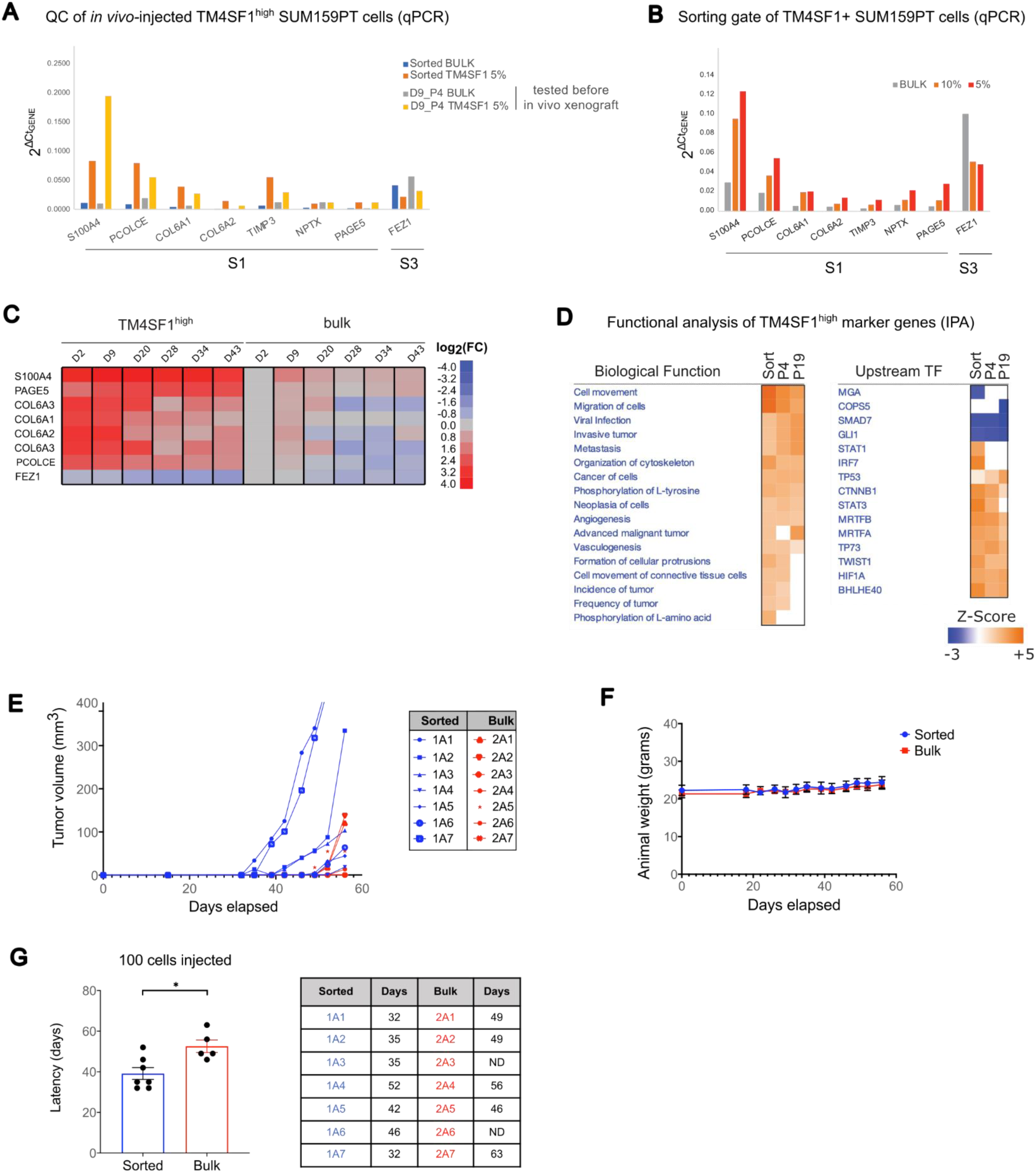
**A.** RT-qPCR data of 7 genes from the S1 gene signature and FEZ1 from the S3 signature respectively in bulk and TM4SF1^high^ cells at time of sorting (day 0) and before in vivo injection (day 9). **B.** RT-qPCR data of the genes shown in **A** at top 5% and top 10% TM4SF1 sorting gates, respectively. **C.** Cell plot showing the expression change (RT-qPCR) of selected genes in the S1 signature in a time-course experiment, where TM4SF1^high^ and bulk cells were let grow unperturbed in 2D for several passages. Entries represent the log_2_(FC) of each gene at every time point normalized over the expression at the first passage (day 2) in bulk cells. **D.** Comparative functional analysis (by Ingenuity Pathway Analysis, IPA) performed on the 195 upregulated genes shown in Figure 2G, indicating the enriched biological functions (left) and upstream transcription factors (right). Entries satisfying |Z-Score| > 2 and p < 0.001 are shown. **E.** Plot showing the growth dynamics of each individual primary tumour derived from inoculation into mammary fat pads of recipient NSG (NOD/SCID/IL2Rγ_c_^-/-^) mice. 100 TM4SF1^high^ cells (at d9) and 100 bulk cells were used, n=7 mice/group. Growth curve after tumours arise until time of euthanisation. **F.** Average weight of mice in response to inoculation with 100 TM4SF1^high^ vs 100 bulk cells in mammary fat pads. Mice weight measurements represent the mean ± SEM, n = 7 mice/group. **G.** Plot showing the latency in days of tumour development for each mouse inoculated with either 100 TM4SF1^high^ or 100 bulk cells. Data are as in **E**. *P = 0.0111; TM4SF1^high^ versus bulk cells, two-sided unpaired t-test. The table on the right reports in detail the latency (days) for each mouse in the two groups, TM4SF1^high^ and bulk.

**Figure S6.**
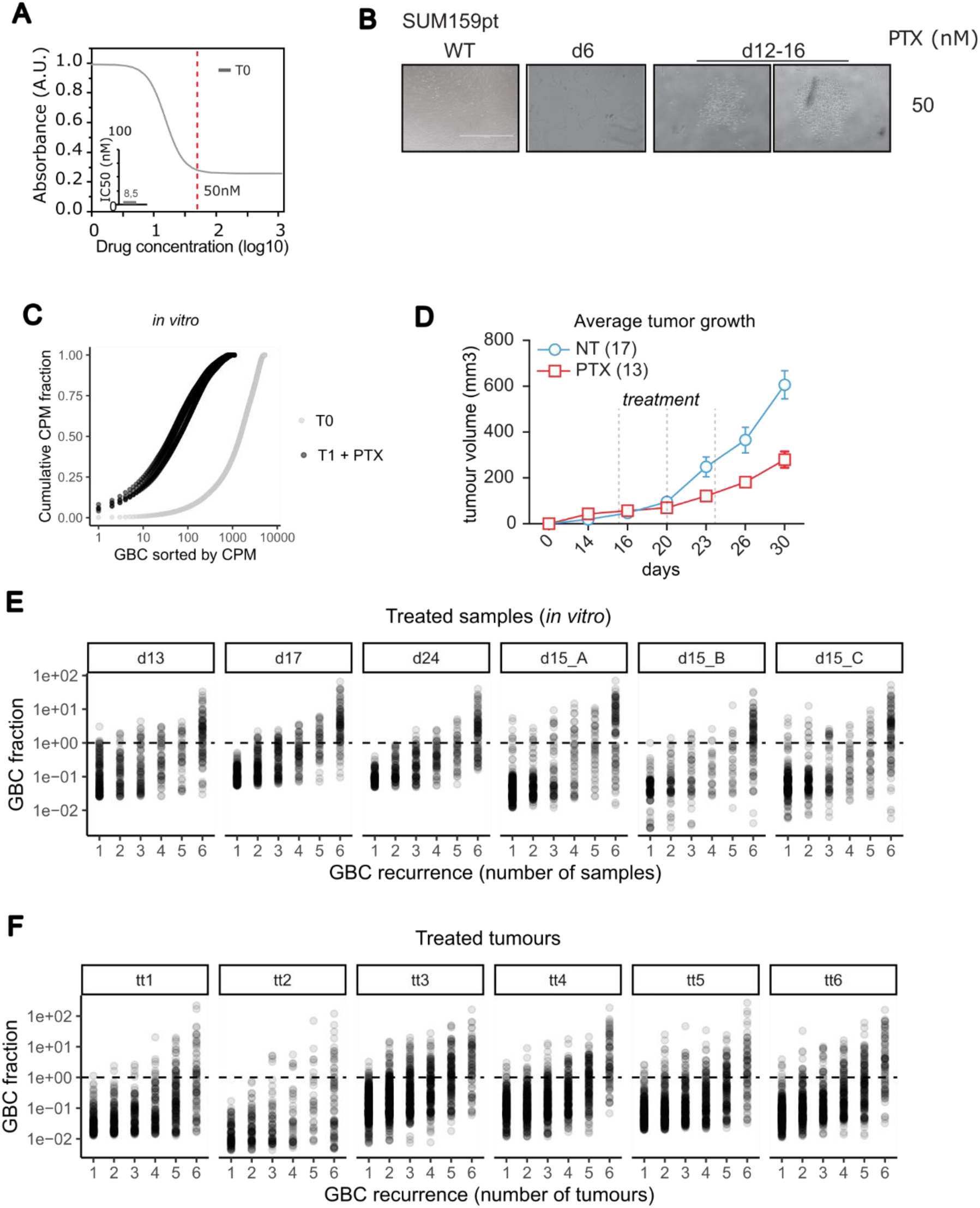
**A.** Dose response curve of SUM159PT treated with paclitaxel (PTX). The curve was estimated according to the Logistic 4P fit model (JMP software). The value of IC50 is reported in the insert. **B.** Representative bright field images of persister colonies that are typically observed 12 days after paclitaxel treatment of SUM159PT cells. Scale bar 1000 uM. **C.** Clone selection upon paclitaxel treatment (50nM) in the independent infection experiment. Comparison of cumulative clone distribution in parental and treated *in vitro* samples. **D.** Growth dynamics of SUM159PT-derived tumours treated intraperitoneally every 5 days with either PTX (10 mg/kg in PBS, 17 mice) or vehicle (PBS, NT, 13 mice). Each point of the growth curves represents the tumour volume expressed as the mean value ± SEM. **E.** Relationship between clone abundance and recurrence in six paclitaxel-treated sample, late time points (day ≥ 13). Each graph refers to a sample and each dot is a clone (GBC); clones are grouped by the number of times they are observed across the six samples. If a clone is such that *x = k* in sample *s*, this means that it is detected in exactly *k* samples (including *s*). The *y* axis is the relative clone abundance in each sample, expressed as the frequency over the total clone count in the sample. **F.** Same as **E** for the six paclitaxel-treated tumours (see also Figure 2C legend).

**Figure S7.**
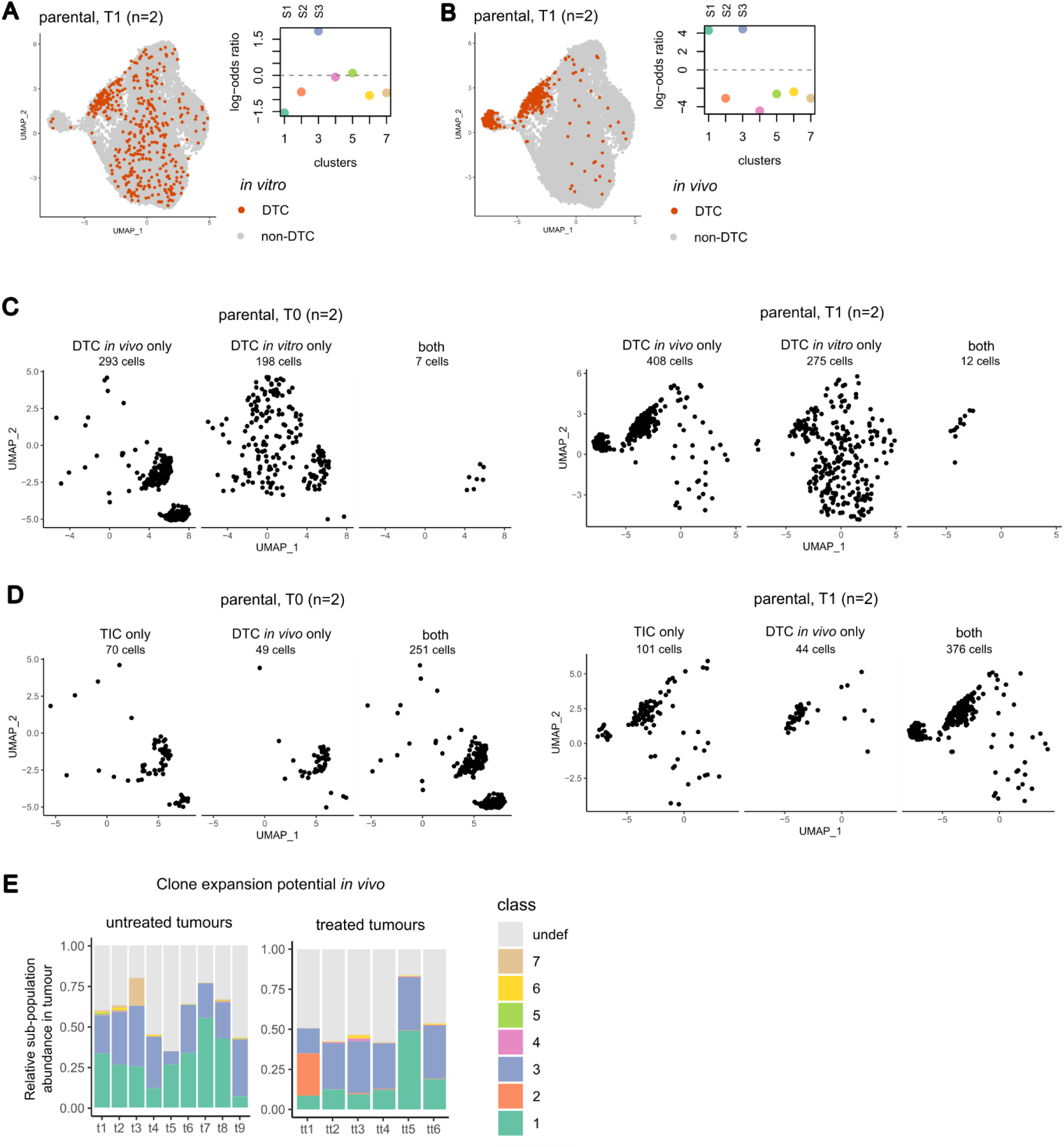
**A.** Mapping of the drug tolerant clones *in vitro* at parental state (T1). Left: UMAP representation of T1 cells on gene expression space; the 287 cells classified as DTC *in vitro* are coloured in orange. Right: the *x* axis is the cluster identifier, and the *y* axis is the log-odds-ratio obtained from the contingency table comparing cluster assignment and DTC labelling *in vitro* across cells at T1. **B.** Mapping of the drug tolerant clones *in vivo* at parental state (T1). Left: UMAP representation of T1 cells on gene expression space; the 420 cells classified as DTC *in vivo* are coloured in orange. Right: the *x* axis is the cluster identifier, and the *y* axis is the log-odds-ratio obtained from the contingency table comparing cluster assignment and DTC labelling *in vivo* across cells at T1. **C.** UMAP representation of T0 (left) or T1 cells (right), split according to whether cells are classified as DTCs only *in vivo*, only *in vitro*, or in both assays. **D.** UMAP representation of T0 (left) or T1 cells (right), split according to whether cells are classified only as TICs, only as DTCs *in vivo*, or in both assays. **E.** Association between parental state (T1) and clone expansion *in vivo*, without and with treatment. Top: bar plot showing the relative abundance of T0 clusters in every untreated (left) or treated tumour (right), respectively (unassigned clusters are shown in grey).

**Figure S8.**
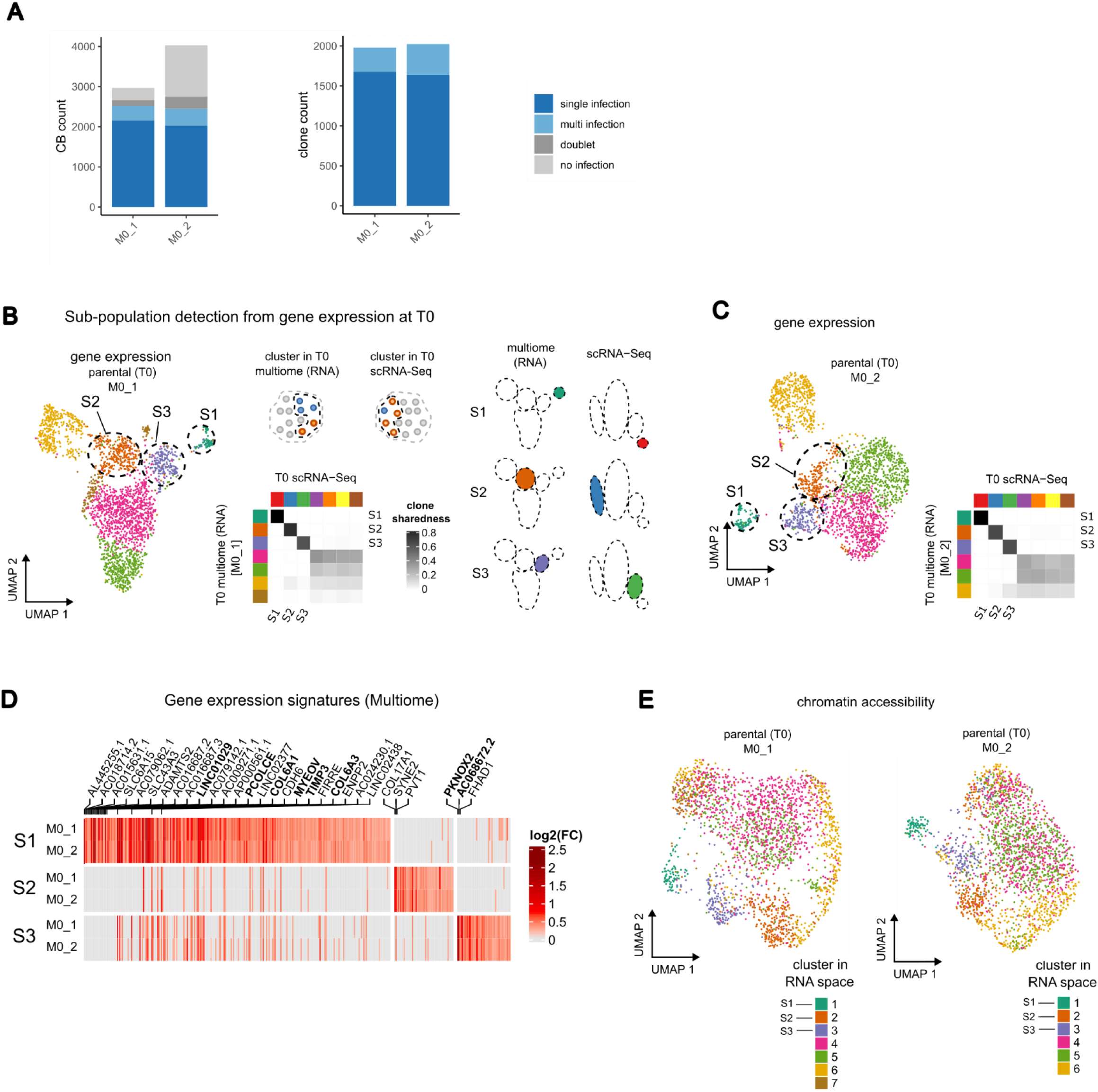
**A.** Clone calling for multiome replicates at T0 (M0_1 and M0_2). Left: cellular barcode (CB) classification into single infection, multi-infection, doublet, and no infection; CB count (left) and clone count (right) are shown (see also **Figure S1B**). **B.** Clone sharedness score between M0_1 and T0. Left: UMAP representation of M0_1 nuclei in gene expression space, coloured by gene expression clusters (2446 nuclei in total). Centre: heatmap where rows are gene expression clusters in M0_1, columns are clusters in T0, and entries are clone sharedness score values for each cluster pair. Rows and columns are sorted according to the pairs with the highest score. The mapping to the subpopulations S1, S2, S3 is indicated for the corresponding clusters in M0_1. Right: schema showing the location of the subpopulations on UMAP. **C.** Clone sharedness score between M0_2 and T0. Left: UMAP representation of M0_2 nuclei in gene expression space, coloured by gene expression clusters (2377 nuclei in total). Right: heatmap where rows are gene expression clusters in M0_2, columns are clusters in T0, and entries are clone sharedness score values for each cluster pair. Rows and columns are sorted according to the pairs with the highest score. The mapping to the subpopulations S1, S2, S3 is indicated for the corresponding clusters in M0_2. **D.** Subpopulation gene signatures in gene expression space (Multiome). Heatmap rows are subpopulations split by sample, columns are genes, and entries are log_2_(FC) values between a subpopulation and its complement within the same sample. All differentially expressed genes (DEGs) in at least one subpopulation are reported (see Methods), the top 20 (S1) or top 5 (S2, S3) are labelled with the corresponding gene symbol, and the ones common to scRNA-Seq signatures are highlighted in bold (see also Figure 1G). **E.** UMAP representation on ATAC space, coloured by gene expression clusters for the 2446 nuclei of M0_1 (left) and the 2377 nuclei for M0_2 (right).

**Figure S9.**
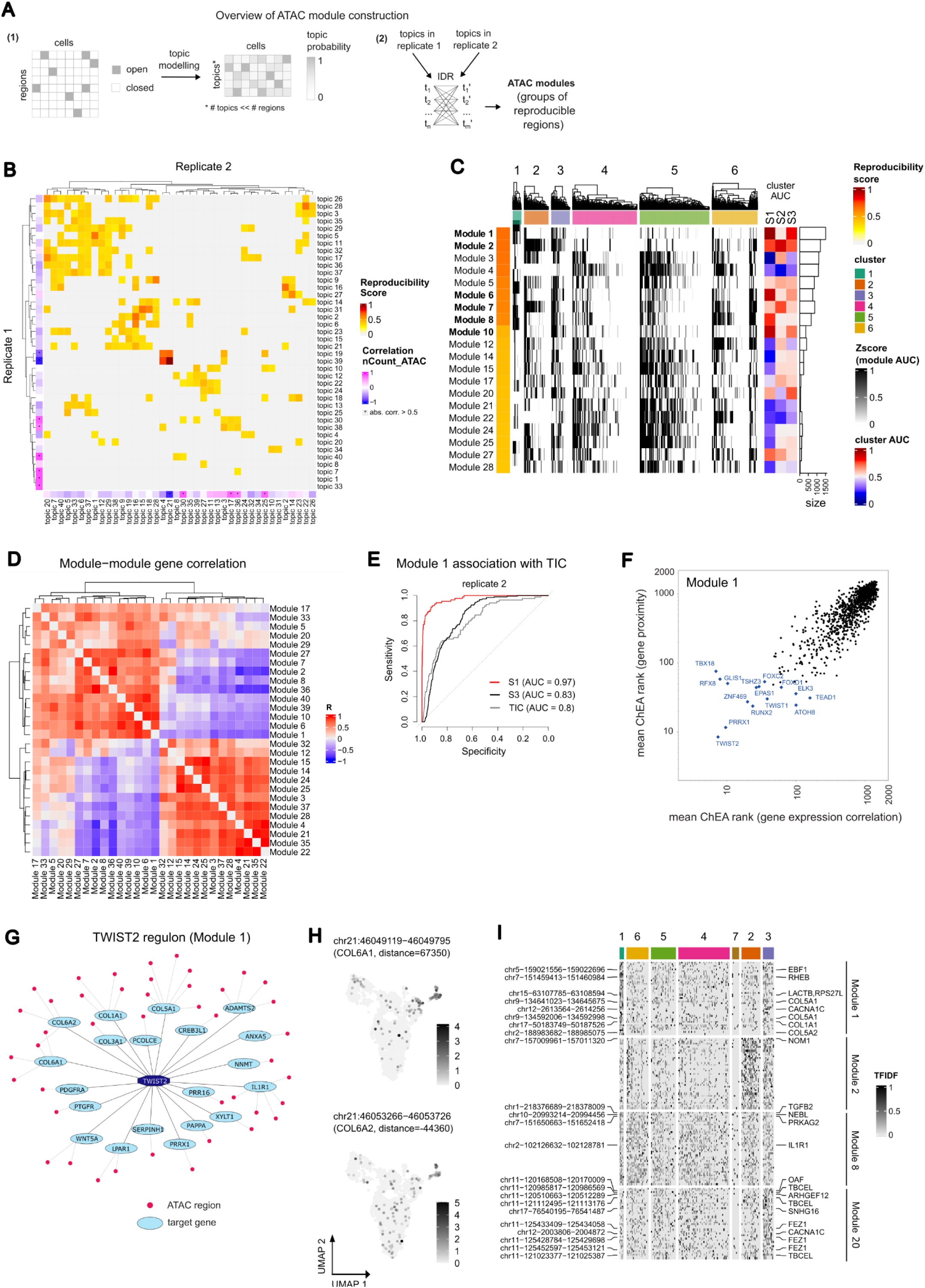
**A.** Schema of the procedure for ATAC module extraction. First, topic modelling is used to extract a small set of topics for each replicate separately (M0_1 and M0_2). A probability value for each (topic,cell) pair is defined, representing the probability of a topic to be present in a cell. A probability value for each (region,topic) pair is also defined, representing the probability of a region in a topic. Then, each topic in M0_1 is compared to each topic in M0_2 using IDR, resulting in a set of regions with low p-value, called *reproducible* regions, for each (topic,topic) pair. These sets of regions are called *ATAC modules*. A reproducibility score is then defined for each ATAC module based on IDR (see Methods). **B.** Topic modelling on ATAC-Seq regions. The heatmap shows the comparison of the output of topic modelling on the two Multiome samples; rows are all topics in replicate 1, columns are all topics in replicate 2, and entries are reproducibility scores (see Methods). Rows and columns are annotated with the Pearson’s R correlation coefficient between (topic,cell) probabilities and ATAC fragment counts across cells. Rows and columns marked with an asterisk are such that |R| > 0.5. **C.** Comparison between ATAC module and gene expression in single nuclei (replicate 2). In the heatmap, rows are the top 20 scoring modules containing ≥ 20 regions, annotated with their size (bar plots on the left), columns are nuclei split by cluster, and entries are module AUC scores representing the overall accessibility of a module in each cell (see Methods). The heatmap annotation on the right shows the degree of association (AUC) of a module (expressed as module AUC score) to subpopulations (S1, S2, and S3), ranging from negative (blue) to positive associations (red). ATAC modules predicting either S1, S2, or S3 with AUC > 0.75 are reported in bold. Hierarchical clustering of columns is performed with `complete` method from `hclust` on Euclidean distances. **D.** Association between ATAC modules by gene expression correlation. Entries represent the Pearson’s R correlation coefficient between gene scores in two ATAC modules; the score is the Spearman’s rho correlation coefficient between gene expression and module AUC score. Here, the top 40 reproducible modules are considered and only the ones with size ≥ 20 are shown. **E.** ROC curves showing the performance of Module 1 AUC as a predictor of S1 (red), S3 (black), or TICs (grey) on Multiome replicate 2. **F.** Transcription factor enrichment analysis obtained with ChEA on genes positively correlated with Module 1 AUC (*x* axis) or genes whose locus is located ≤ 100 kbp away from any region in Module 1 (*y* axis). Values report the average TF rank obtained with ChEA on the two gene sets, respectively. The top 10 ranked genes for either gene set are labelled. **G.** The “TWIST2 regulon” includes the set of genes whose expression is positively correlated with Module 1 AUC and that i) are predicted as TWIST2 targets by ChEA3 and ii) possess a proximal Module 1 ATAC-region (<100 kbp from the gene). **H.** UMAP plot showing the term frequency-inverse document frequency (TFIDF) score for the two selected target regions of Module 1 in M0_1. **I.** Regions in modules and gene locus proximity. The top 40 reproducible regions (IDR score) for 4 modules are reported (module labelling as in Figure 4D). Rows are regions and columns are nuclei split by cluster. Hierarchical clustering of columns is performed with `complete` method on Euclidean distances. Entries are term frequency-inverse document frequency (TFIDF) scores. Regions located at ≤ 50 kbp from any gene in subpopulation gene signatures (obtained from scRNA-Seq samples only, see Figure 1F) are labelled.

**Figure S10.**
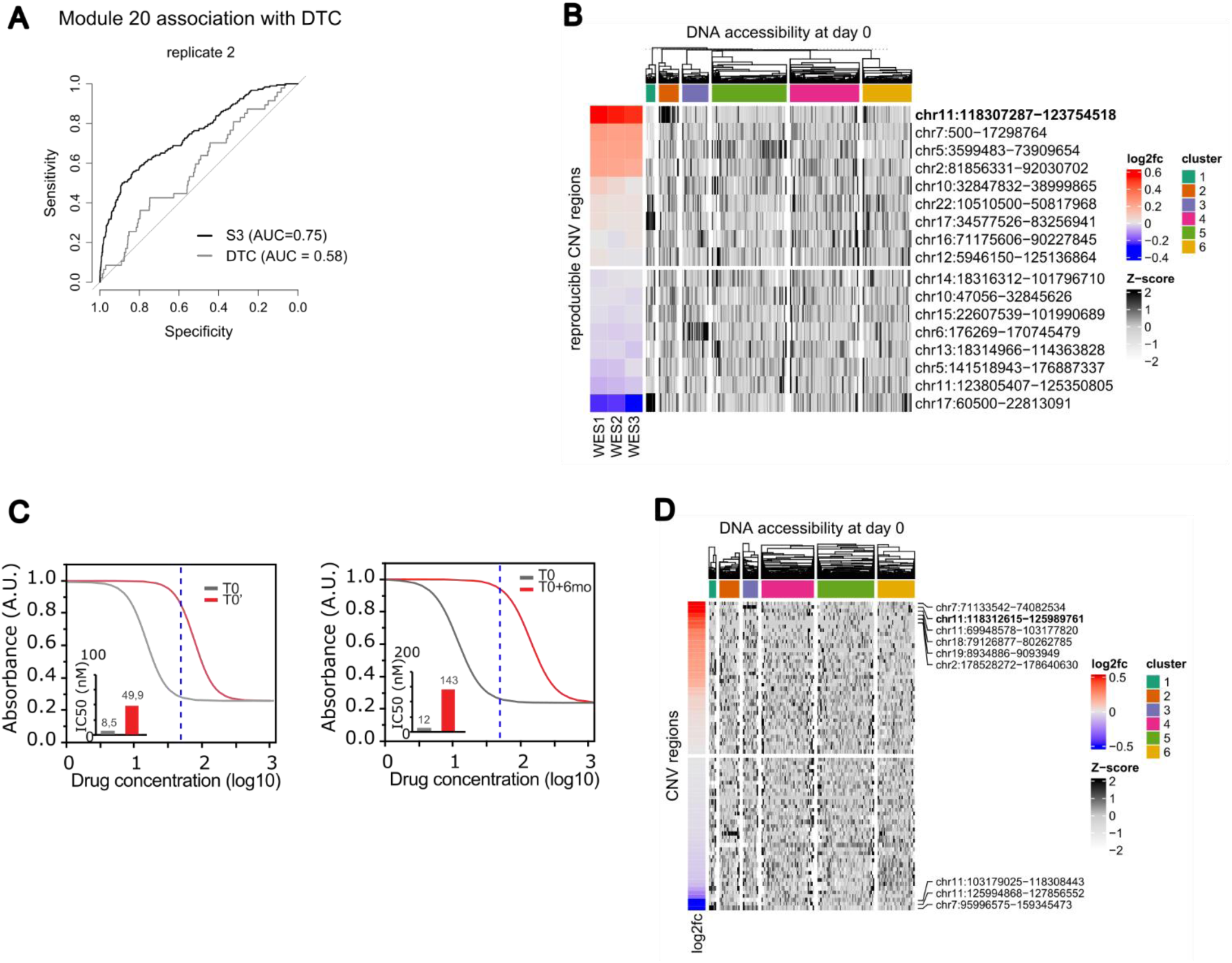
**A.** Association between Module 20 and S3 subpopulation (replicate 2). Top: ROC curve showing the power of Module 20 AUC score to predict (i) the membership to the DTC in vitro pool (grey line) or (ii) the membership to the S3 subpopulation (black line) on replicate 2. **B.** Copy-number variants (CNVs) pre-and post-treatment (drug tolerance assay, Multiome replicate 2; see Figure 5D). Heatmap showing the association between ATAC-Seq signal from Multiome data and copy-number variants inferred by WES. Rows are consensus CNV obtained from the analysis of paclitaxel-treated samples (day 15; n=3 replicates), as in the drug-tolerance experimental design *in vitro* from Figure 3A; columns are nuclei in Multiome replicate 2; entries are cumulative ATAC counts in each CNV locus and in each nucleus. Rows correspond to consensus CNVs across three replicates. The coverage log_2_(FC) between each treated sample and the untreated reference (baseline SUM159PT cells) is reported on the left; the chromosome location is shown on the right. Columns are grouped by gene expression cluster; hierarchical clustering is performed with `complete` method from `hclust` on Euclidean distances. **C.** Dose response curve of different SUM159 populations (T0, T0’ and T0+6months, as defined in Figure 5D-E) treated with paclitaxel. The curve) was estimated according to the Logistic 4P fit model (JMP software). IC50 values are reported in the insert. **D.** CNVs pre- and post-treatment (drug resistance assay, Multiome replicate 2; see Figure 5E). The heatmap shows the association between ATAC-Seq signal from Multiome data (replicate 2) and copy-number variants predicted by WES (see also A). Rows are CNVs obtained from the analysis of paclitaxel-treated samples (n=1; see Figure 5E).

**Figure S11.**
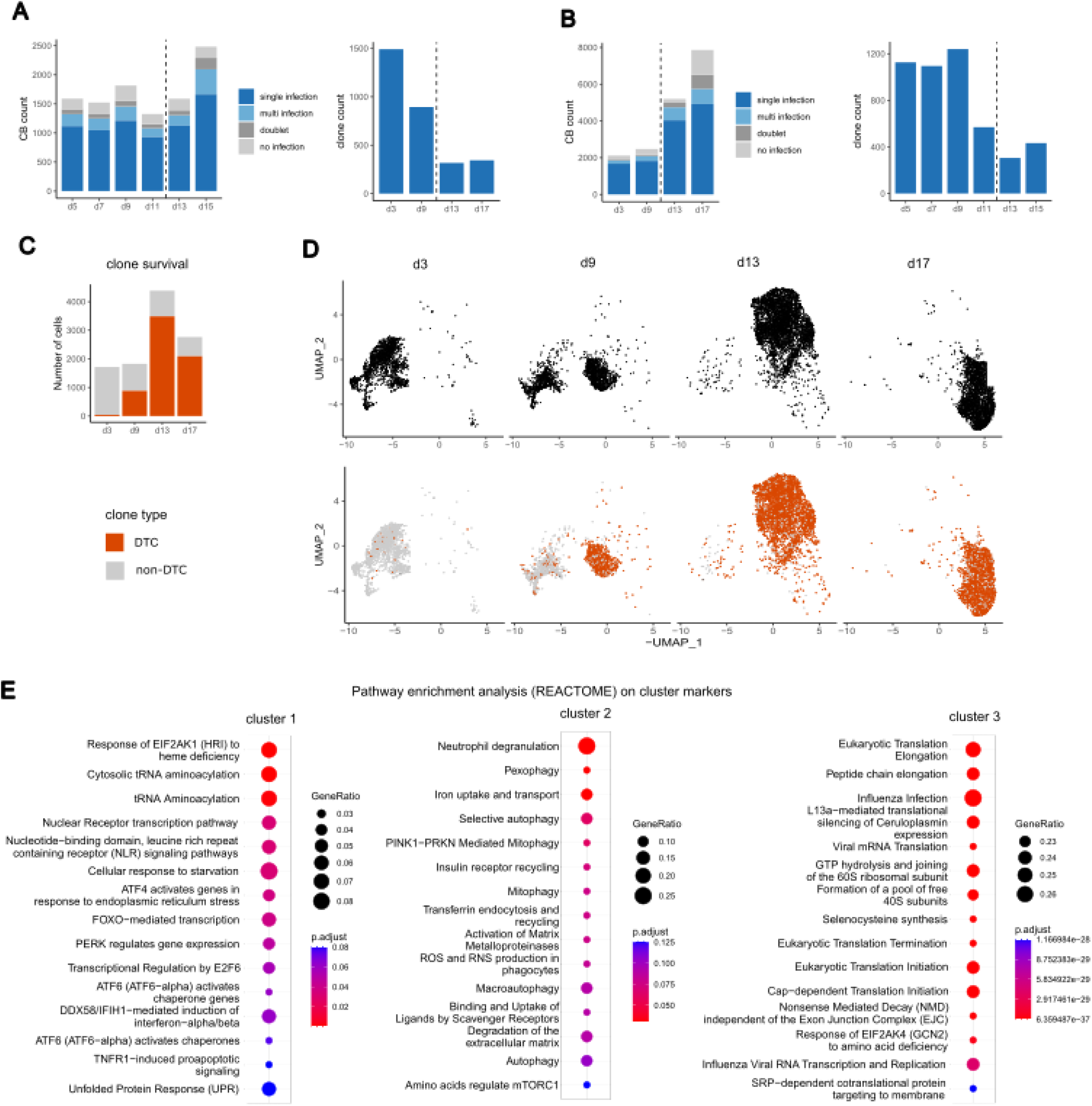
**A.** Clone calling for time-course drug tolerance assay (replicate 1). Left: cellular barcode classification into single infection, multi-infection, doublet, and no infection; CB count (left) and clone count (right) are shown (see also **Figure S1B**). **B.** Clone calling for time-course drug tolerance assay (replicate 2). See **A**. **C.** Bar plot showing the distribution of DTC *in vitro* on treated samples (replicate 2). **D.** Drug-tolerant clone selection across time (replicate 2). Cells from treated samples are mapped to a common gene expression UMAP space (10698 cells in total), split by sample (top), coloured depending on whether they are drug tolerant (in orange, as defined in **C**) or not (in grey). **E.** Pathway enrichment analysis (REACTOME) of significantly up-regulated genes in clusters of drug-treated samples (as in Figure 6C-D). The top 15 significantly enriched terms (q-value < 0.1) are reported and sorted by non-increasing q-value. The size of the circles is proportional to the fraction of genes found in each pathway for each subpopulation signature.

**Figure S12.**
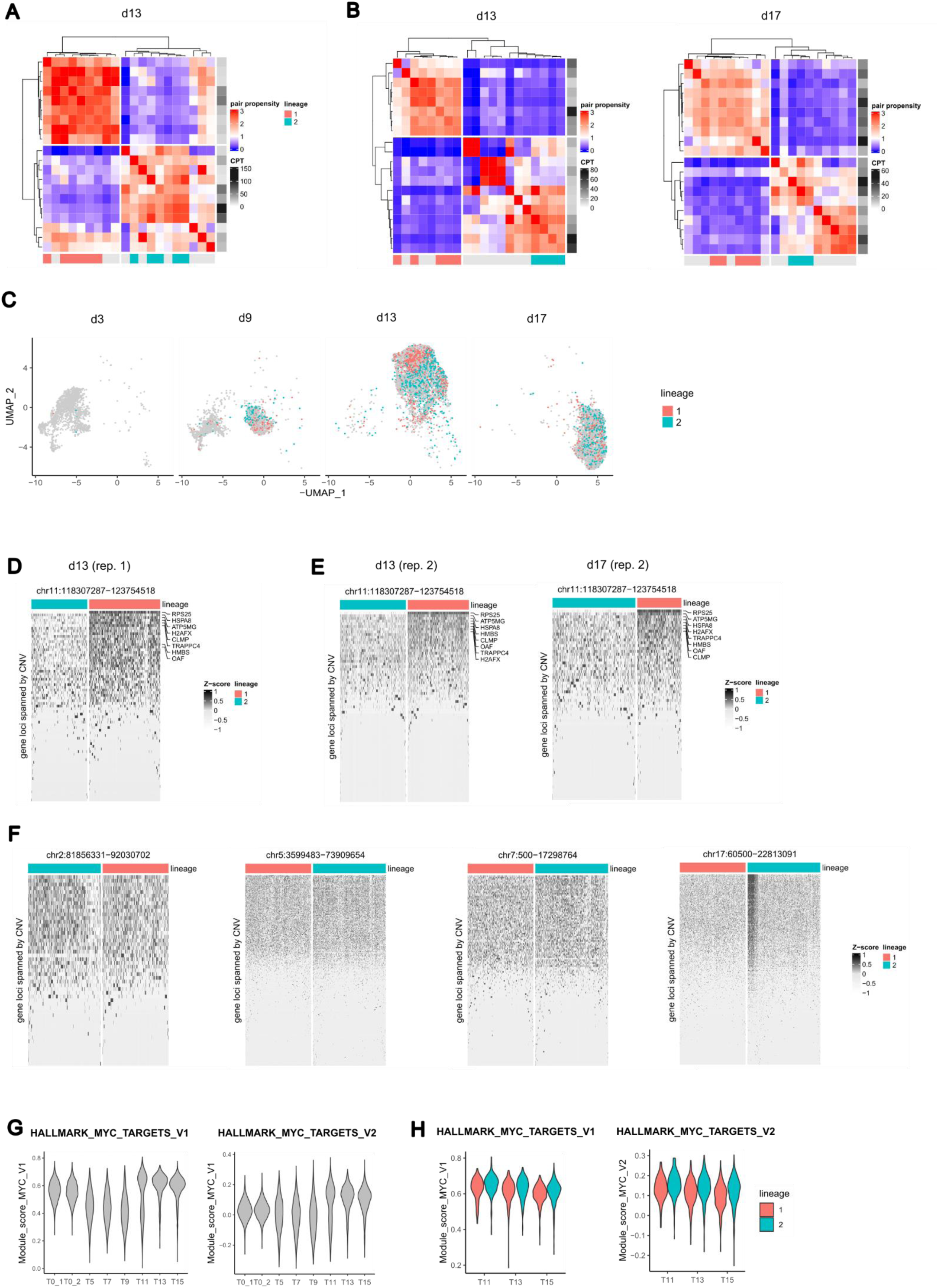
**A.** Pair propensity value of top expanded clone pairs at day 13 (exp1; clones *i* with *p_ii_* < 1 are not shown); rows and columns are distinct clones, rows are annotated by clone abundance (in CPT, count per thousand cells), and columns are annotated by lineage. **B.** Pair propensity value for top expanded clone pairs at day 13 and 17 (exp2). See also **A**. **C.** UMAP representation of cells at day 3, 9, 13 and 17 (exp2) and coloured according to lineage assignment (lineage 1 in pink, lineage 2 in cyan, and unassigned clones in grey). **D.** Heatmap where rows are gene loci spanned by the highest log_2_(FC) consensus amplification in drug treated samples, which is located on chromosome 11 (see Figure 5A), columns are cells at day 13 assigned to lineages (exp2), and entries are log-normalised, scaled UMI counts. Rows are sorted by non-increasing average expression. Columns are split by lineage and clustered with `complete` method on Euclidean distances. **E.** Heatmap relative to chromosome 11 amplification, where columns are cells at day 17 (replicate 2). See also **D**. **F.** Heatmaps relative to all the other consensus CNVs such that average |log_2_(FC)| > 0.1, for day 15 (replicate 1). See also **D**. **G.** Quantification of MYC activity. Distribution of MYC activity scores across all time points, before (T0, see Figure 1B) and after treatment (day ≥ 5, see Figure 6A); MYC activity is computed using the Hallmark signatures of MYC Targets (V1 and V2) from The Molecular Signatures Database (**MSigDB**) with ModuleScore function. **H.** Distribution of MYC activity score, as in **F**, at day ≥ 11 and split by lineage.

## Supplementary table legends

**Suppl. Table 1.** Cell annotation for scRNA-Seq and Multiome datasets (RNA-Seq library only).

**Suppl. Table 2.** Differential expression analysis at baseline (scRNA-Seq). DEA between each cluster and its complementary at baseline in the same experiment (n = 2 each).

**Suppl. Table 3.** Functional annotation of differentially expressed genes at baseline. DEA between each cluster and its complementary. REACTOME database is used for functional annotation.

**Suppl. Table 4.** Clone detection in vitro and in vivo from bulk GBC sequencing. Count statistics and classification of GBC sequences.

**Suppl. Table 5.** Differential expression analysis of SUM159 derived tumours compared to parental cells. Expression level in TPM, L2FC between TM4SF1-high and bulk and DESeq2 statistics.

**Suppl. Table 6.** Analysis of [Pal et al.] dataset. Epithelial cells from four TNBC samples are reported.

**Suppl. Table 7.** Differential expression analysis of TM4SF1-high cells compared to bulk population. Expression level in TPM, log_2_(FC) between TM4SF1-high and bulk and edgeR statistics.

**Suppl. Table 8.** Differential expression analysis at baseline (Multiome). DEA between each cluster and its complementary at baseline in the same experiment (n = 2 each).

**Suppl. Table 9.** CisTopic analysis and ATAC module detection. CisTopic is run for the two Multiome replicates separately. IDR is computed for each cross-replicate topic pair.

**Suppl. Table 10.** Spearman’s rho between module AUC (Module 1) and gene expression across cells, for the two replicates separately.

**Suppl. Table 11.** Transcription factor enrichment by ChEA3. Meanrank score and overlapping genes using either Module 1 proximal genes (GREAT) or Module 1 correlated genes (Spearman’s rho) as query.

**Suppl. Table 12.** Copy number variants called on Whole Exome Sequencing using CNVkit. Paclitaxel treated samples at day 15 (D15A-B-C) or day 90 are compared with parental cells.

**Suppl. Table 13.** Clone detection from single-cell RNA-Seq. Count statistics and classification of clones.

**Suppl. Table 14.** Differential expression analysis after treatment. DEA between lineage 1 and lineage 2 after treatment in the same experiment (all time points). DEA between each cluster and its complementary after treatment in exp2.

**Suppl. Table 15.** Functional annotation of differentially expressed genes after treatment. DEA between each cluster and its complementary. REACTOME database is used for functional annotation.

**Suppl. Table 16.** RT-qPCR oligos, sgRNAs sequences, primers pair for GBC PCR on gDNA, primers pair for GBC PCR (DROP-seq).

